# Functional profiling of the *Toxoplasma* genome during acute mouse infection

**DOI:** 10.1101/2023.03.05.531216

**Authors:** Christopher J. Giuliano, Kenneth J. Wei, Faye M. Harling, Benjamin S. Waldman, Madeline A. Farringer, Elizabeth A. Boydston, Tammy C. T. Lan, Raina W. Thomas, Alice L. Herneisen, Allen G. Sanderlin, Isabelle Coppens, Jeffrey D. Dvorin, Sebastian Lourido

## Abstract

Within a host, pathogens encounter a diverse and changing landscape of cell types, nutrients, and immune responses. Examining host-pathogen interactions in animal models can therefore reveal aspects of infection absent from cell culture. We use CRISPR-based screens to functionally profile the entire genome of the model apicomplexan parasite *Toxoplasma gondii* during mouse infection. Barcoded gRNAs were used to track mutant parasite lineages, enabling detection of bottlenecks and mapping of population structures. We uncovered over 300 genes that modulate parasite fitness in mice with previously unknown roles in infection. These candidates span multiple axes of host-parasite interaction, including determinants of tropism, host organelle remodeling, and metabolic rewiring. We mechanistically characterized three novel candidates, including GTP cyclohydrolase I, against which a small-molecule inhibitor could be repurposed as an antiparasitic compound. This compound exhibited antiparasitic activity against *T. gondii* and *Plasmodium falciparum,* the most lethal agent of malaria. Taken together, we present the first complete survey of an apicomplexan genome during infection of an animal host, and point to novel interfaces of host-parasite interaction that may offer new avenues for treatment.

## INTRODUCTION

Pathogens encounter complex and dynamic environments within the body of a host. To survive and replicate, pathogens must navigate diverse tissues and their constituent cell types, innate and adaptive immune responses, and restricted or fluctuating nutrient abundances. Despite their utility, cell culture models of microbial pathogenesis fail to capture many of the specific challenges pathogens face within their hosts. Probing biological processes in their organismal context demands the development of new methods. Recent advances have enabled studies of complex processes directly in mice, highlighting novel aspects of cell-cell communication^1^, embryology^2^, stem cell differentiation^3^, and spatially-restricted transcriptional responses^4^. Motivated by this framework, we developed approaches to characterize the genome-wide fitness landscape of the model apicomplexan parasite *Toxoplasma gondii* during infection of a natural animal host.

Apicomplexans comprise a phylum of obligate intracellular parasites that includes the etiological agents of human diseases such as malaria (*Plasmodium* spp.), cryptosporidiosis (*Cryptosporidium* spp.), and toxoplasmosis (*T. gondii*). Despite the significant burden of infection associated with these pathogens, many aspects of their biology remain poorly defined. *T. gondii* is often used as a model of the apicomplexan phylum because of its ease of culturing and genetic manipulation. *T. gondii* is also widespread, infecting over 25% of the global population^5^ and a vast number of warm-blooded species^6^. Mice are among the natural intermediate hosts of *T. gondii*, making them an ecologically-relevant model of infection. These combined experimental traits make *T. gondii* ideally suited to study host-parasite interactions of a human-infective pathogen.

Genetic screens remain one of the most powerful tools to probe the molecular basis of biological processes, yet implementing them in a whole organism poses significant technical challenges. Despite this, screens in animals have effectively captured phenotypes that are inaccessible in cell culture, informing aspects of bacterial infection^7^, liver development^8^, and cancer metastasis^9^. The adaptation of CRISPR/Cas9 screening in *T. gondii* has enabled unbiased, broad-based examination of genetic contributions to parasite fitness^10^. By varying the selective environments, screens have exposed gene function in relation to specific pressures, such as drug treatment^11^, oxidative stress^12^, and innate immunity^13^. To date, however, genome-wide surveys of parasites have been largely restricted to cell culture, failing to capture factors that address challenges these pathogens only encounter within an animal host.

Efforts to screen parasites in animal hosts have been limited. Recent small-scale screens of *T. gondii* in mice have revealed new determinants of infection^14–16;^ however, due to difficulties in maintaining library coverage, these screens only targeted 149 to 235 genes at a time— representing only ∼2.5% of the over 8,000 genes in the parasite genome. Vector insertion–based screens in *Plasmodium berghei* individually disrupted 2,578 genes across 58 pools as parasites replicated in mice^17^. Although this work surveyed ∼50% of the genome, the inability to culture *P. berghei* outside of mice precludes direct comparison of fitness contributions between cell culture and whole-organism environments.

To address the challenges of screening parasites in mice, we designed a condensed-barcoded gRNA library. Barcoded CRISPR screens have been shown to decrease experimental noise in a variety of low-efficiency screening contexts^18–21^. Adding to these methodologies, we employ a population genetics-based analysis that is broadly applicable across experimental systems where bottlenecks limit screening efficiency. Leveraging these methods, we present the first genome-wide survey of an apicomplexan during infection of an animal host.

## RESULTS

### Generation of condensed and barcoded gRNA libraries for screening in mice

Previous attempts to screen in mice were hampered by the difficulty of maintaining gRNA library coverage during mouse infection^14–16^. To address this challenge, we tested whether more condensed genome-wide libraries would perform well in cell culture. We designed a new genome-wide library following current guidelines for gRNA efficiency^22^, cloning libraries with either nine or three gRNAs per gene—the latter representing a subset of gRNAs from the larger library (**Table S1**). We conducted screens using these two libraries in cell culture and calculated gRNA scores as the fold change of a gRNA’s final abundance relative to the initial library. Comparing the mean gRNA scores for each gene (fitness score) indicates the results of the three-gRNA library are consistent with those of the nine-gRNA version, as well as previously published genome-wide screens (**Fig. 1A, S1A–B, Table S2**).

**Figure 1.**
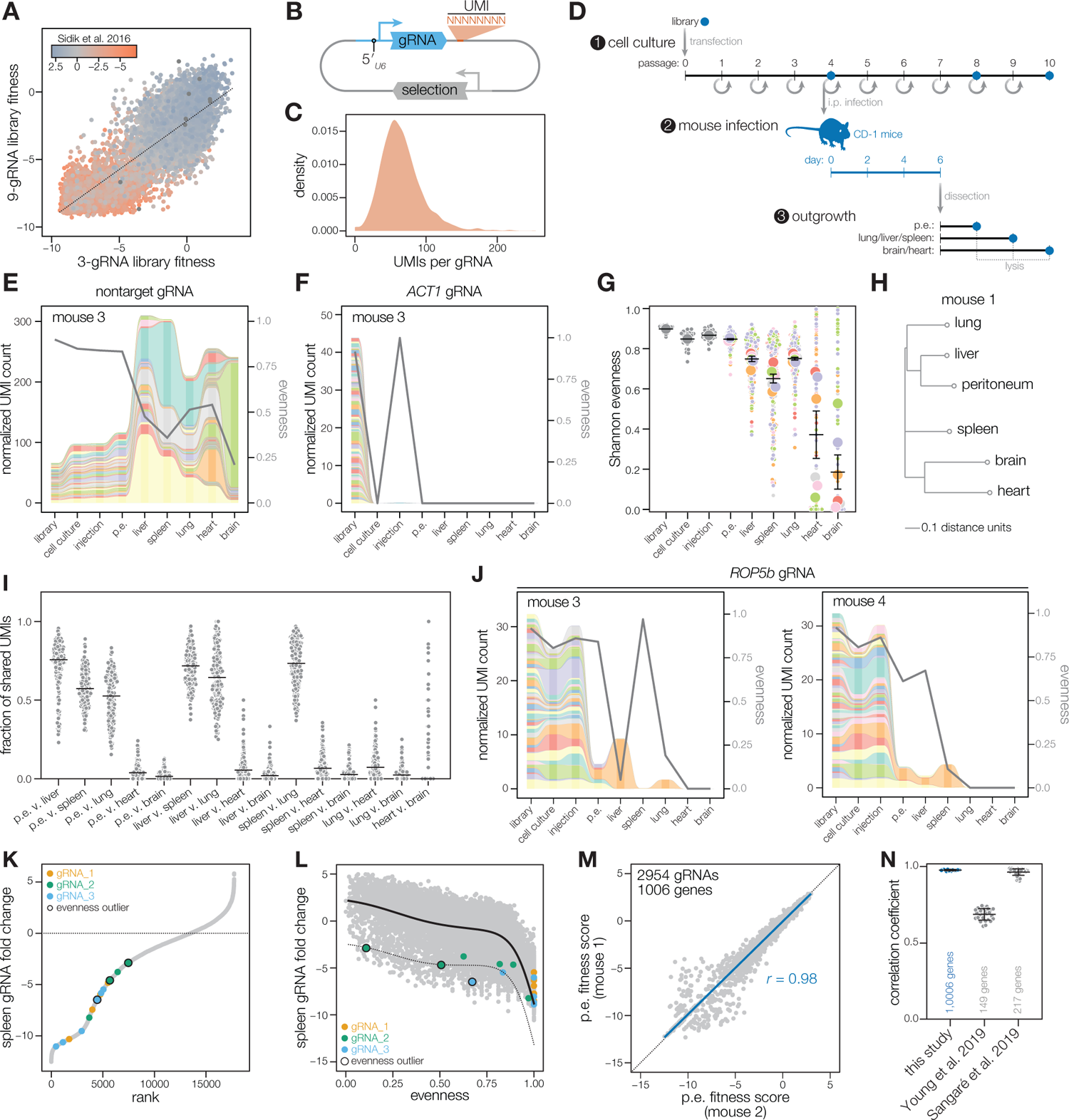
Condensed-barcoded gRNA libraries improve screening efficiency in mice. **(A)** Scatter plot of gene fitness scores generated from CRISPR screens with nine-gRNA and three-gRNA libraries. Data points are colored based on published phenotype scores from Sidik, *et. al* 2016^10^. **(B)** Design of barcoded gRNA constructs. **(C)** Density curve of barcodes per gRNA. **(D)** Screening workflow. **(E–F)** Alluvial plots depicting relative barcode abundance of a nontargeting gRNA **(E)** or an *ACT1*-targeting gRNA **(F)** across tissues in a representative mouse. Line graph indicates evenness of barcodes in each sample. **(G)** Superplot of Shannon evenness for nontargeting gRNAs across tissues from six mice. Colors indicate different mice with small points for each gRNA and large points for the mean of all gRNAs in each mouse. Mean ± SEM for all mice is overplotted. **(H)** Dendrograms of tissue similarity based on genetic distance of barcode relatedness from nontargeting gRNAs. **(I)** Jaccard similarity (fraction of shared UMIs for a given nontargeting gRNA) between indicated tissue pairs across six mice. Black bars indicate mean fraction. **(J)** Alluvial plots depicting relative barcode abundance of a *ROP5b* (*TGGT1_411430*) gRNA across tissues in two mice. Line graphs indicate evenness of barcodes in each sample. **(K)** Rank-ordered spleen gRNA fitness scores from six mice. **(L)** Spleen gRNA fitness scores plotted against evenness. Blue lines represent thresholds for gRNA outliers at two standard deviations from the generated model. **(M)** Correlation of gRNA scores between two replicate mice (Pearson’s correlation *r* = 0.98). **(N)** Correlations among replicate mice in this study compared to previously published small-scale *T. gondii* mouse screens. Mean ± SD indicated.

Further, we used unique molecular identifiers (UMIs) to track clones derived from independent integrations of the same gRNA. The UMIs are sequences of eight random base pairs included in our gRNA vectors, such that each gRNA in the library is tagged with 65 ± 29 (SD) barcodes (**Fig. 1B–C**). By tracking barcoded parasites, we can identify the number of independent clones that contribute to a gRNA’s relative abundance. A gRNA with multiple UMIs implies that many clones contribute to the observed phenotype. Conversely, high relative abundance of a gRNA represented by a single UMI could indicate random jackpotting through bottlenecking or reflect failed disruption of the targeted gene.

To limit the number of parasites needed to maintain library coverage, we divided the *T. gondii* genome into 17 libraries that could be screened independently (**Table S1**). Each library included control gRNAs that target genes known to be fitness-conferring in cell culture or required during mouse infection, and a panel of 30 nontargeting gRNAs.

### Barcoded parasite populations track dissemination dynamics

As a pilot study, we combined two libraries to screen 1,006 genes in a single experiment. We transfected Cas9-expressing parasites with our gRNA library and maintained the population in culture for four passages. During the fourth passage, a portion of the parasite population was used to infect outbred CD-1 mice intraperitoneally (i.p.), while continuing to passage the remainder of the population in cell culture for the duration of the experiment. Six days after infection, parasites were harvested from infected mice by isolating peritoneal exudates (p.e.), liver, spleen, lung, heart, and brain. Parasites from each sample were expanded briefly in cell culture before isolating genomic DNA (**Fig. 1D**). We found that a brief period of outgrowth was optimal for scoring gene fitness compared to directly processing infected tissues, as this would reduce DNA contributed from dead parasites (**Fig. S1C, Table S3**). The outgrowth period differed between tissues, with p.e. samples typically lysing the monolayers in approximately two days; lung, liver, and spleen in approximately four days; and heart and brain in approximately six days. The outgrowth period is consistent with the predicted parasite burdens of the various tissues following i.p. infection^14, 23^.

We examined the dynamics of infection by analyzing barcodes of parasites that received nontargeting gRNAs. Nontargeting gRNAs were generally represented by many UMIs, although the diversity of the populations was clearly diminished in tissues distal from the injection site (**Fig. 1E, S1D, Table S4**). By contrast, gRNAs targeting the essential gene Actin (*ACT1*, *TGGT1_209030*)^24, 25^ were quickly depleted, with all clones cleared from the population in all tissues (**Fig. 1F, S1E**). We used metrics from population genetics to quantify these dynamics: species richness counts the number of distinct clones found in a sample, while Shannon evenness quantifies how evenly represented each clone is in the population. These metrics indicated that population diversity is largely maintained in tissues with high parasite burdens like the peritoneum, liver, spleen, and lung, but is significantly reduced due to bottlenecks in the heart and brain (**Fig. 1G, S1F**).

We next investigated whether parasites follow a reproducible path of dissemination or instead seed distal tissues, such as the brain, directly from the peritoneum. By comparing barcode frequencies, we calculated the genetic distance between parasite populations from different organs and constructed dendrograms to represent their relatedness (**Fig. 1H, 1SG–H**). Parasite populations in distant tissues like heart and brain were often the most distinct, with distance metrics approximately 1.7-fold greater than those between other tissues (**Fig. S1I**). Indeed, many of the UMIs present in the heart and brain are absent from tissues more proximal to the injection site, indicating that parasites can seed the heart and brain directly from the peritoneum (**Fig. 1I**).

Population analyses can be compiled for each gene targeted in our screen, enabling us to identify abnormalities in gRNA scores arising from bottlenecks and jackpotting. For example, the *ROP5* locus contains a cluster of polymorphic alleles known to be dispensable in human fibroblasts but required during mouse infection^26–28^. In most mice, gRNAs targeting *ROP5b* (*TGGT1_411430,* **Table S1**) were consistently depleted from all tissues. However, in some samples, such as the liver of mouse 3 and the spleen of mouse 4, a single *ROP5b* gRNA failed to drop out due to the persistence of a dominant clone (**Fig. 1J**). The occurrence of the same dominant clone in different mice likely indicates an ineffective mutagenesis event, wherein Cas9 failed to generate a loss-of-function mutation in *ROP5b*.

We can systematically identify instances of bottlenecks and ineffective mutagenesis by measuring the evenness of UMIs for each gRNA. However, we found evenness scores produced a variable distribution where high fitness scores tended toward lower evenness. Neutral gRNAs therefore exhibit high relative abundance yet low evenness due to the difficulty of maintaining large numbers of neutral clones. By contrast, gRNAs targeting essential genes have low fitness scores yet high evenness, as it is easier to maintain low diversity in small populations. To identify outlier gRNAs, we established a dynamic threshold using a Bayesian machine learning approach, which selects gRNAs that deviate from these trends in an unbiased manner. For example, this filter removed the *ROP5b* gRNA 2 from the spleen of mouse 4, increasing the consistency of fitness scores among all *ROP5b* gRNAs in this tissue across mice. Altogether, we excluded 3,107 gRNA values from the individual tissue samples analyzed (2.83% of all gRNA values; **Fig. 1K–L**). Using this approach, we were able reproducibly screen over 1000 genes in a single mouse, improving the efficiency of screening in mice by an order of magnitude over previous studies (**Fig. 1M–N**)^14–16^.

### Genome-wide contributions to parasite fitness during mouse infection

We deployed the improved methods to screen the 17 sublibraries that comprise the entire parasite genome. Five of these sublibraries were designed around specific sets of genes, such as the presence of a signal peptide, localization to secretory organelles (rhoptries, dense granules, or micronemes), or predicted metabolic function (**Fig. 2A, Table S1**). The remainder of the genome was randomly divided so that each library targeted approximately 500 genes. The sublibraries were screened in pairs, culturing the transfected parasite population for four passages before splitting it to infect mice or continue passaging in culture (**Fig. 1D**). Parasites were recovered through outgrowth from tissues harvested six days post infection. For each subscreen, we sequenced the gRNAs from the original library, the injected population (passage four), outgrowth samples, and the continued passages in cell culture (passages eight and ten). The final dataset comprised relative gRNA abundances from 329 samples originating from the nine screens across 54 mice.

**Figure 2.**
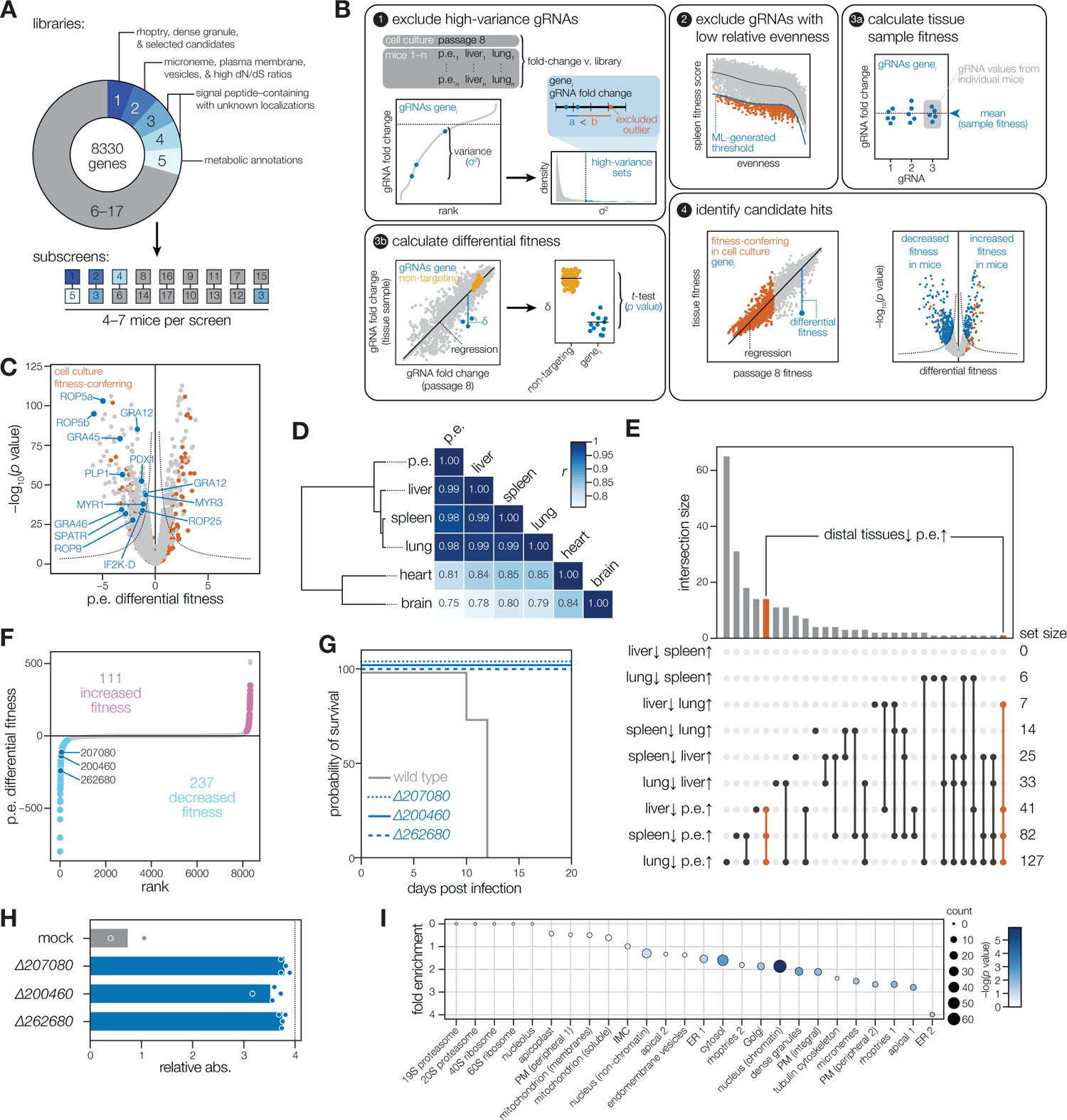
Genome-wide survey of gene fitness during mouse infection. **(A)** Partitioning of the genome over 17 sublibraries screened across nine subscreens. **(B)** Analysis pipeline used to calculate differential gene fitness scores between tissues and cell culture. **(C)** Volcano plot showing differential fitness score from parasites harvested from mouse peritoneum compared to cell culture (unpaired two-tailed *t*-test). Dotted line indicates a threshold of [-log_10_(pval) · differential fitness score] = 30. Genes required for fitness in cell culture are highlighted in red. Genes referenced in the text as specifically required for fitness in mice are highlighted in blue. **(D)** Heatmap of Pearson correlations (*r*) of gene fitness scores between indicated tissues. **(E)** Upset plot of candidates with differential fitness in indicated tissue comparisons. In each comparison, candidates are scored with decreased fitness in the first tissue relative to the second one. **(F)** Rank-ordered plot of differential fitness between mouse peritoneum and cell culture, highlighting candidates that surpass a threshold of [-log_10_(pval) · differential fitness score] = 30. **(G)** Survival of CD-1 mice after infection with 10,000 parasites of the indicated genotype. **(H)** Serum response to *T. gondii* antigens from surviving mice. Dotted line indicates maximum limit of detection. **(I)** Hypergeometric gene enrichment analysis of 348 candidates with differential phenotypes in mice.

We computed gene fitness scores as the average fold change in gRNA abundance for gRNAs targeting a given gene. We first filtered out outlier gRNAs from the top 5% most variable gene sets, as these may indicate gRNAs with consistently poor on-target or off-target activity (558, or 2.1% of all gRNAs). We next analyzed barcode dynamics for each gene, and applied our evenness filter to exclude likely instances of bottlenecks and ineffective mutagenesis (23,652 values, 2.59% of whole dataset; **Fig. 2B**). Control gRNAs behaved consistently across subscreens (**Fig. S2A–D, Table S5**). Additionally, replicate screens of one of the sublibraries proved highly correlated, allowing us to compare fitness scores of genes from different subscreens (Pearson’s correlation, *r* = 0.943; **Fig. S2E**).

We compared fitness scores from each mouse tissue to those derived from cell culture to identify genes that contribute to parasite fitness during the acute stages of mouse infection. To identify top candidates, we computed a linear regression between fitness scores from cell culture and a given mouse tissue, and determined a distance metric for each gRNA relative to the linear model (**Fig. 2B, Table S5**). For a given gene, gRNA distances across mice were averaged for each tissue type, generating tissue-specific gene fitness scores. Genes with cell culture fitness scores below –5 were excluded to focus our analysis on genes that primarily influence fitness during mouse infection.

Comparisons between fitness scores in cell culture and the peritoneum identified several previously characterized factors required for mouse infection (**Fig. 2C**). Some of these factors influence parasite-intrinsic processes that impact fitness in mice by altering gene expression, metabolic adaptation, invasion, or egress. For example, the eukaryotic initiation factor 2α kinase (*TgIF2K-D*; *TGGT1_319610*), found to promote extracellular viability in cell culture^29^, was necessary for fitness in mice, indicating that parasites may experience increased extracellular stress during mouse infection. We similarly captured various metabolic genes required for mouse infection, highlighting differences in nutrient availability. For example, we identified *PDX1* (*TGGT1_237140*), a gene involved in the biosynthesis of the ubiquitous cofactor vitamin B6. Loss of *PDX1* is compensated by nutrient availability in cell culture but renders parasites avirulent in mice^30, 31^. We also captured *SPATR* (*TGGT1_293900*) and *PLP1* (*TGGT1_204130*), which facilitate efficient parasite invasion and egress, respectively; both are required during mouse infection, yet become dispensable in cell culture due to the extended times afforded to parasites to invade and egress^32–34^. Our results suggest that under the stringent conditions experienced during mouse infection, parasites require greater efficiency in diverse cellular processes.

Our screens also captured factors that may contribute to host modulation and immune evasion. In pooled screens, loss of secreted effectors may be complemented by coinfection of host cells with wildtype and mutant parasites. Indeed, such trans-complementation has been shown to mask phenotypes of known effectors in previously published small-scale screens^15^. Nevertheless, we detected several secreted effectors important during mouse infection. We recapitulated the requirement for the *ROP5a* (*TGGT1_308090*) and *ROP5b* (*TGGT1_411430*), virulence factors that suppress host immunity-related GTPases (IRGs) that disrupt the parasitophorous vacuole^26, 27^. We identified additional secreted effectors that lack detailed functional characterization, including *ROP9* (*TGGT1_309590*), *ROP25* (*TGGT1_202780*), *GRA12* (*TGGT1_288650*), *GRA25* (*TGGT1_290700*), and *GRA46* (*TGGT1_208370*)^35–40^. Beyond individual virulence factors, we captured three known components of the secretion machinery that transports effectors across the parasitophorous vacuole membrane into the host cytosol, including *MYR1* (*TGGT1_254470*), *MYR3* (*TGGT1_237230*), and *GRA45* (*TGGT1_316250*)^13, 41–43^. This indicates that our pooled screening approach challenges parasites to compete independently, suggesting that novel secreted virulence factors are among the candidates with decreased fitness in mice.

### Tissue-specific effects on parasite fitness

*Toxoplasma* disseminates systemically^44^, raising the possibility that parasites require particular adaptations to infect and survive in different tissues. We compared fitness scores between anatomical sites in search of tissue-specific phenotypes. Fitness scores were highly correlated between tissues, indicating that parasites experience similar selective pressures in these environments during the six-day infection period captured by our screen (**Fig. 2D**). Heart and brain samples were less correlated to other organs, consistent with the strong bottlenecks experienced by the parasite population in these tissues (**Fig. 1G**). Such bottlenecks also reduced the reproducibility of scores from heart and brain samples across mice (**Fig. S2F**). We therefore focused further analyses of tissue-specific effects on comparisons between the p.e., liver, spleen, and lung samples. We performed a regression-based analysis to compare fitness scores between these tissues, and identified 220 genes with tissue-specific phenotypes (**Fig. 2E, Table S6**).

The lung yielded the greatest number of tissue-specific candidates. Candidate divergence in the lung mirrors its anatomical location outside of the peritoneum, in contrast to the liver and spleen. Lung samples display lower species richness compared to liver or spleen, which may increase the false-positive rate of tissue-specific phenotypes (**Fig. 1G**). Consistent depletion from multiple tissues increases confidence in certain candidates. Comparing the fitness scores from the liver, spleen, and lung to those from the peritoneum revealed 15 candidates with consistently diminished fitness in distal tissues (**Fig. 2E, S2G–I**). Some of these candidates initially displayed decreased fitness in the p.e. relative to cell culture; however, these phenotypes were exacerbated as the infection disseminated to other organs. Mutants in *PDX1* and *PDX2*, two genes involved in the production of vitamin B6 precursors^30^, were initially depleted from the peritoneum, but their relative abundance dropped further in distal tissues. This may imply an increase in certain metabolic demands as parasites disseminate. By contrast, other candidates displayed moderate fitness scores in the peritoneum yet dropped out in distal organs. Mutants in *ROM4* (*TGGT1_268590*), a gene implicated in efficient parasite motility^45^, were significantly depleted in distal tissues relative to the peritoneum. These examples illustrate the range of fitness profiles captured by the screens.

Our results highlight the challenge of unequivocally identifying tissue-specific effects in mouse screens. A dissemination phenotype was previously reported for *TgWIP* (*TGGT1_247520*)^14^, but this effect was not detected in our screens (**Fig. S3A–D**). To further verify our results, we regenerated a *TgWIP* knockout and did not observe a dissemination phenotype (**Fig. S3E–F**). We conclude that the effects of TgWIP may be context dependent. In aggregate, many tissue-specific effects appear to result from additive fitness costs as parasites disseminate from the peritoneum. Nevertheless, our results suggest that intermediate fitness defects compound differently for certain genes during the course of infection.

### Mutants with enhanced fitness in mice

Comparison between growth in cell culture and the peritoneum provided the most robust measure of fitness in mice, based on the population diversity of these samples (**Fig. 1G**). Genes with differential fitness effects in the peritoneum were rank-ordered based on the magnitude and statistical significance of their divergence from a linear model of their effects in cell culture. 348 genes displayed significantly different fitness during mouse infection compared to cell culture (**Fig. 2F**). Of note, 111 genes in this list are candidates for which loss appears to promote increased fitness in mice.

Among the genes whose loss improved parasite fitness in the peritoneum relative to cell culture were a putative copper-transporting P-type ATPase (*TgCuTP*; *TGGT1_201150*), a thioredoxin reductase (*TgTr*; *TGGT1_309730*), and the glycosyltransferase *GNT1* (*TGGT1_315885*). TgCuTP localizes to the plant-like vacuole and acidocalcisomes, and is suspected to function in copper homeostasis. While TgCuTP has not been functionally characterized in *Toxoplasma*, disruption of *PbCuTP* in *Plasmodium berghei* reduced parasite fitness in cell culture but not during mouse infection^46^. This suggests there is increased pressure on copper regulation in cell culture compared to mouse infection. Analogously, differences in oxidative stress may underlie the effects of *TgTr* and *GNT1.* TgTr is a conserved enzyme involved in reactive oxygen species (ROS) scavenging. While *ΔTgTr* parasites exhibit growth defects both in cell culture and during mouse infection^12, 47^, our results indicate that the defects in mice are less severe. GNT1 belongs to a conserved pathway that regulates Skp1 through glycosylation, and has been associated with O_2_ sensing in *Toxoplasma* and *Dictyostelium*^48, 49^. GNT1 was upregulated during adaptation to cell culture, and its loss reduced parasite proliferation^48, 50^. Levels of atmospheric oxygen (21%) used in cell culture, in excess of the physiological range (2–9%), may increase the demand for O_2_ sensing and response pathways^51^. These examples illustrate that replication in mice may compensate for fitness defects apparent in cell culture, as is the case for *CuTP* and *TgTr*. Alternatively, some of the identified mutants may authentically improve parasite fitness in mice.

### Identification of genes that contribute to parasite virulence

Our analysis revealed 237 genes that selectively support parasite fitness during mouse infection (**Fig. 2F**). Mutation of 17 of these genes has been previously shown to attenuate parasite virulence in mice (**Table S7**). An additional 15 genes have not been directly studied in mice, but their loss impacts growth under specific culture conditions that may capture aspects of mouse infection, such as growth in interferon gamma–stimulated macrophages^13, 29, 42^. To validate candidates with decreased fitness in mice, we generated knockout strains for three previously uncharacterized genes by replacing their coding sequences with a green-fluorescent marker (**Fig. S4A**). Two candidates, *TGGT1_262680* and *TGGT1_200460*, lack any identifiable domains. TGGT1_200460 was recently localized to the Golgi^52^, whereas the localization of TGGT1_262680 is unknown. The third candidate, *TgMYST-B* (*TGGT1_207080*), participates in histone acetylation and the DNA-damage response^53^. As predicted by our screen results, all three knockout strains were avirulent in mice, despite viability in cell culture (**Fig. 2G–H, S4B**). The predicted function of TgMYST-B suggests that parasites experience greater strains on genome stability during mouse infection, possibly due to the increase in reactive oxygen species associated with inflammation^54, 55^. Our screens similarly recapitulated previously characterized decreased virulence of other DNA repair mutants including DNA damage inducible protein 1 (*TgDDI1: TGGT1_304680*) and *Rad23* (*TgRad23: TGGT1_295340*), further emphasizing the importance of these pathways during animal infection^13, 56^.

Since parasite fitness is context dependent, we considered how species differences may influence the outcome of our screens. Following standard practice, we use human fibroblasts for routine culture of *Toxoplasma*. To address whether some of the effects observed in mice reflect host-species differences, we conducted genome-wide screens during infection of two immortalized mouse embryonic fibroblast (MEF) cell lines, and compared fitness scores to our screen results in human fibroblasts. Fitness scores were highly correlated between the different host cell contexts. Only six of the 348 candidates with differential fitness in mice scored as significant in comparisons of either MEF line to human fibroblasts (**Fig. S4C–F, Table S8**). This indicates that most of these candidates are not arising from species differences alone, but are rather derived from the pressures of the organismal environment.

Our validation efforts provide further support for the over 300 newly identified candidate genes influencing parasite fitness in mice. This gene set was enriched for candidates likely to interact with the host (localized to secretory organelles such as rhoptries, micronemes, and dense granules) as well as possible regulation of parasite gene expression (nuclear chromatin localizations)^52^. By contrast, gene categories likely to be essential in cell culture, such as localization to the mitochondrion, apicoplast, nucleolus, and the proteasome, were not enriched among our candidates (**Fig. 2I**). To further understand the demands on parasite fitness experienced during mouse infection, we characterized three additional candidates (*TGGT1_235130, TGGT1_272460*, and *TGGT1_253780*) in greater detail.

### RASP1 facilitates repeated invasion and broadens host cell tropism

We investigated candidates localized to rhoptries, given critical functions played by these organelles in infection and rewiring of host cells^57^. Among the top-scoring mouse dependencies was the rhoptry-localized *RASP1* (*TGGT1_235130*) (**Fig. 3A**). During preparation of this manuscript, a small-scale CRISPR screen also found *RASP1* to be important for mouse infection^16^. *RASP1* and its paralog *RASP2* have been described based on co-expression with known rhoptry genes^58^. Both Rhoptry Apical Surface Proteins (RASPs) localized to the cytosolic face of the rhoptries. While RASP2 was essential for parasite invasion and rhoptry protein secretion, *Δrasp1* parasites behaved normally in cell culture. We confirmed that RASP1 co-localizes with a rhoptry marker to the parasite apical end (**Fig. 3B, S5A**). Homologous proteins in *Plasmodium falciparum*, PfCERLI1 and PfCERLI2, have been similarly localized, and individually contribute to merozoite invasion of erythrocytes^59, 60^. Notably, these studies do not explain why *RASP1* is necessary in mice.

**Figure 3.**
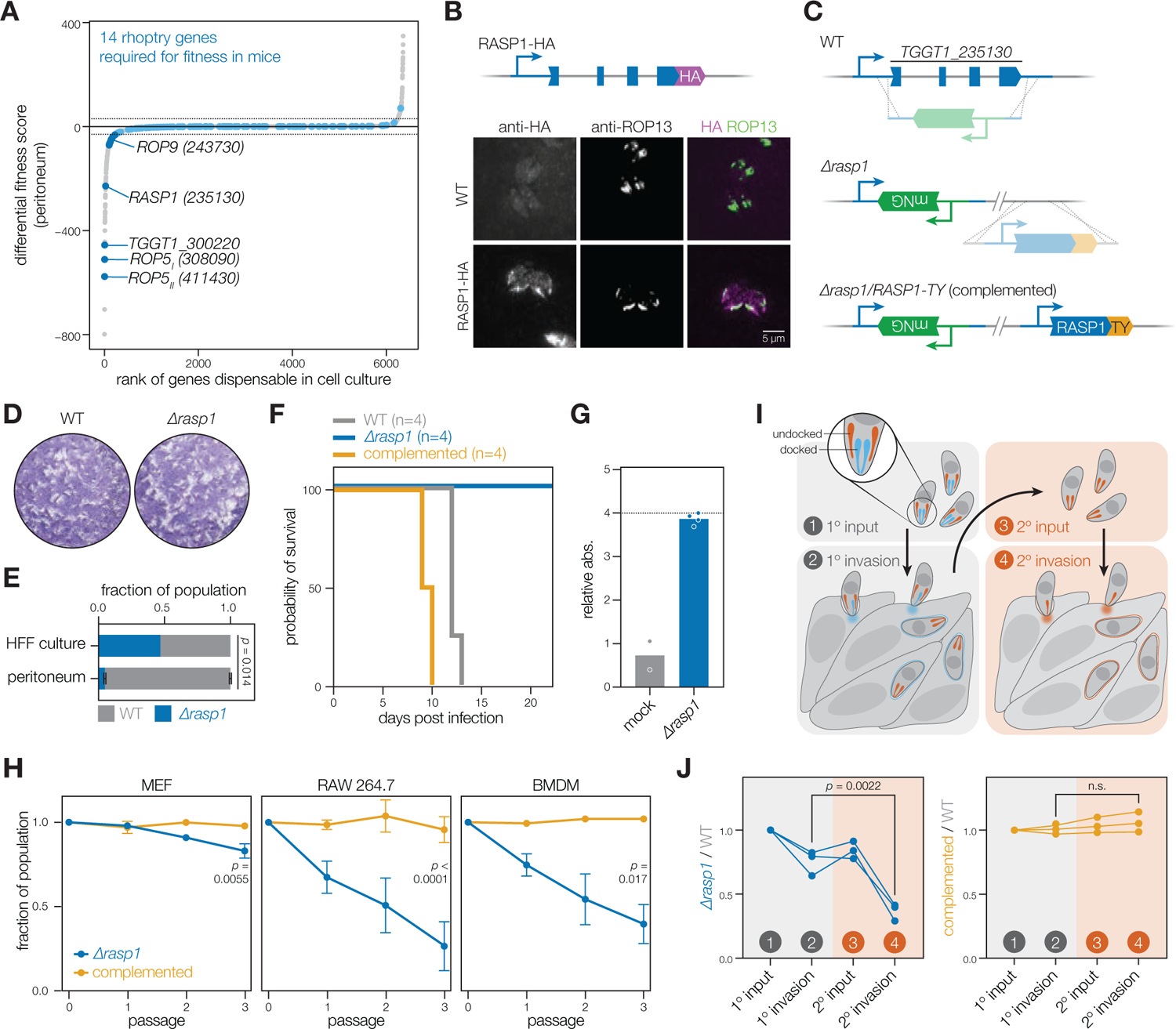
RASP1 facilitates repeated invasion and broadens host cell tropism. **(A)** Differential fitness scores of rhoptry-localized genes among all genes dispensable in cell culture. **(B)** Immunofluorescence images showing the localization of endogenously HA-tagged RASP1. WT parasites are included for reference. **(C)** Schematic of *RASP1* knockout and complementation. **(D)** Plaque assay of *Δrasp1* and WT parasites. **(E)** Competition of *Δrasp1* and WT parasites in cell culture and mouse infection. Mean ± SD indicated (*n* = 2 mice, unpaired two-tailed *t*-test). **(F)** Survival of CD-1 mice after infection with 10,000 WT, *Δrasp1*, or complemented parasites. Number of mice in each group (*n*) is indicated. **(G)** Serum response to *T. gondii* antigens from surviving mice. Dotted line indicates maximum limit of detection. **(H)** Competitions of *Δrasp1* or complemented parasites against WT, in MEFs, RAW264.7, or CD-1 bone-marrow derived macrophages. (*n* = 3 for MEF and RAW264.7, *n* = 2 for BMDM, unpaired two-tailed *t*-test of passage 3 fractions). **(I)** Diagram of the repeated invasion assay, highlighting the various samples measured. **(J)** Relative fraction of *Δrasp1* or complemented parasites during competition with a WT strain in the repeated invasion assays (*n* = 3, ratio paired two-tailed *t*-test).

To study RASP1’s function, we generated knockout parasites by replacing the coding sequence with an mNeonGreen fluorophore (*Δrasp1*; **Fig. 3C, S5B**). We generated a complemented strain by expressing a TY epitope–tagged RASP1 from a safe-harbor locus in *Δrasp1* parasites^61^. The complementing allele was also apically localized (**Fig. 3C, S5C**). Consistent with RASP1 being dispensable in cell culture, we found that *Δrasp1* parasites formed normal plaques in human fibroblast monolayers (**Fig. 3D**). To validate the importance of RASP1 during mouse infection, we performed competitions of WT and *Δrasp1* parasites in cell culture and mouse infection. *Δrasp1* parasites were highly depleted from the peritoneum of infected mice, confirming RASP1 is fitness-conferring during mouse infection (**Fig. 3E**). As competitions only measure relative fitness, we next directly assessed the virulence of *Δrasp1* parasites in mice. *Δrasp1* parasites resulted in no fatalities despite seroconversion, while RASP1-TY complementation restored virulence (**Fig. 3F–G**).

Parasites encounter a variety of host cell types when establishing a successful infection in mice. Replication in macrophages is critical during the early stages of infection, based on their abundance in the peritoneum and role disseminating parasites to distal tissues^62–64^. We therefore compared the fitness of mutant parasites during growth in fibroblasts and macrophages. *Δrasp1* parasites showed a negligible fitness defect in fibroblasts, but were consistently outcompeted in bone marrow-derived macrophages and in the immortalized RAW264.7 macrophage cell line (**Fig. 3H**). Our results suggest a cell type-specific dependency on RASP1 that may drive depletion of *Δrasp1* parasites from the mouse peritoneum.

Previous reports found that the RASP1 paralog RASP2 is necessary for rhoptry secretion^58^, raising the possibility that RASP1 may support rhoptry secretion in a context-specific manner. Only two of the 8–12 rhoptries in a *T. gondii* tachyzoite are docked at the apical end and primed for secretion at a given time^65^. A single rhoptry-secretion event is considered sufficient for host-cell invasion as invasive stages of the related parasite *Cryptosporidium parvum* harbor only a single rhoptry^66^. *T. gondii* appears capable of secreting rhoptry-localized effectors into host cells that are not ultimately invaded^67^. This phenomenon requires that parasites reload their docked rhoptries to enable repeated rounds of discharge. We therefore wondered whether RASP1 participates in rhoptry reloading. Using a modified invasion assay, we measured the ability of *Δrasp1* parasites to undertake multiple rounds of invasion as a proxy for sequential rounds of rhoptry discharge. In these experiments, a mixed population of WT and *Δrasp1* parasites was allowed to invade fibroblasts. Soon after this primary invasion, we mechanically released the invaded parasites and assayed their ability to invade new host cells (**Fig. 3I**). *Δrasp1* parasites invaded host cells at near WT rates during primary invasion, but significantly dropped out when challenged to an immediate second round of invasion—a defect rescued by complementation (**Fig. 3J**). These results suggest that RASP1 is required for repeated invasion attempts, likely by mediating rhoptry reloading.

### GRA72 is required for parasitophorous vacuole maintenance and development

Parasites secrete numerous proteins from dense granules that are necessary for parasitophorous-vacuole (PV) maintenance and host-cell modulation. Our screen captured several previously characterized dense granule protein genes (GRAs), that decreased fitness during mouse infection when disrupted (**Fig. 4A**). The top-scoring candidate from this set was *TGGT1_272460*, a gene of unknown function localized to dense granules by spatial proteomics^52^. An endogenous HA-epitope tag showed TGGT1_272460 is secreted from parasites, and colocalizes with the PV-resident protein MAG1^68^ (**Fig. 4B, S6A**). Based on the observed localization, we designated TGGT1_272460 as a bonafide dense granule protein, naming it *GRA72* per convention.

**Figure 4.**
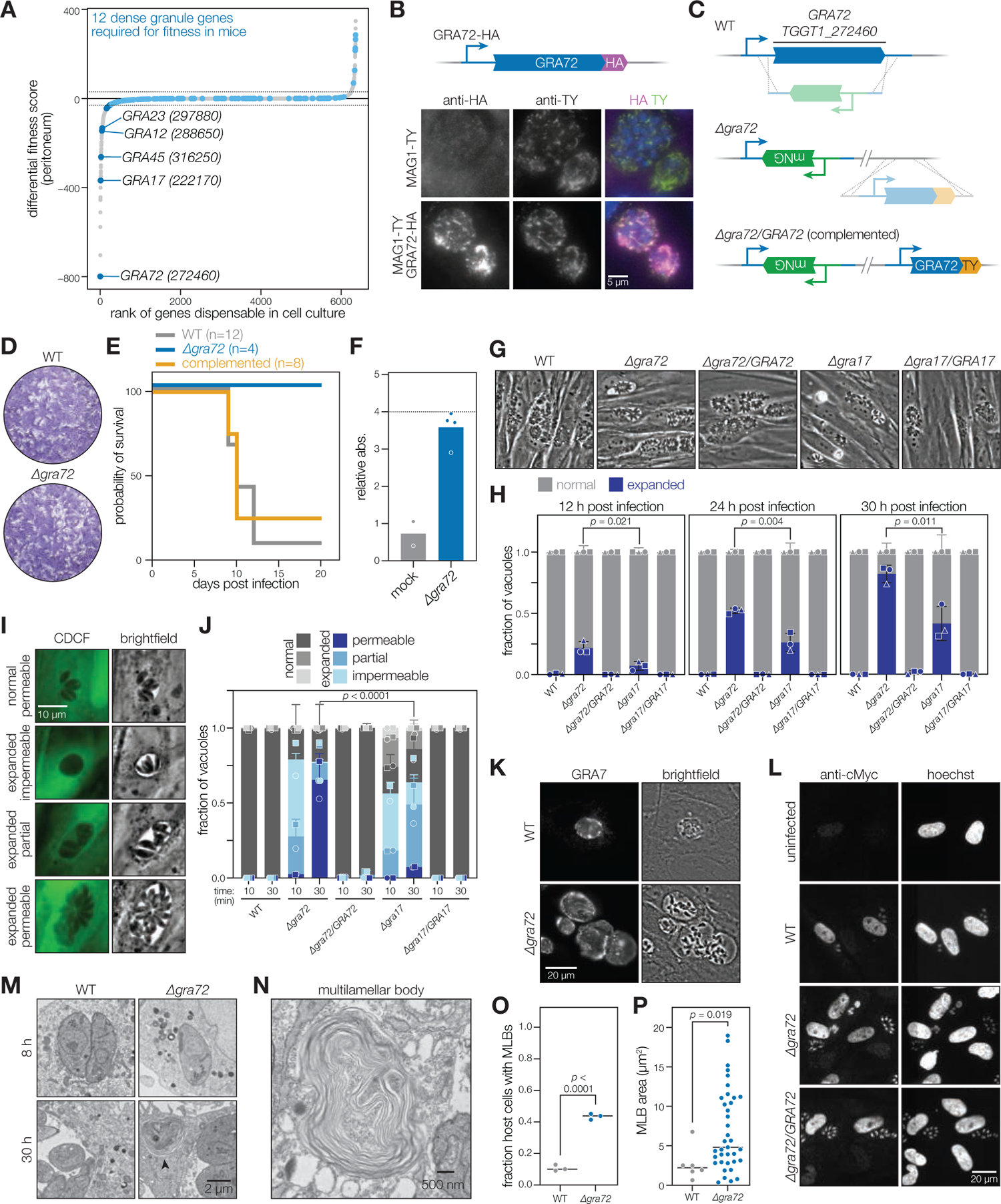
GRA72 is required for parasitophorous vacuole maintenance and development. **(A)** Differential fitness scores of dense granule-localized genes among all genes dispensable in cell culture. **(B)** Immunofluorescence images showing the localization of endogenously HA-tagged GRA72 and endogenously TY-tagged MAG1. **(C)** Schematic of *GRA72* knockout and complementation. **(D)** Plaque assay of *Δgra72* and WT parasites. **(E)** Survival of CD-1 mice after infection with 10,000 WT, *Δgra72*, or complemented parasites. Number of mice in each group (n) is indicated. **(F)** Serum response to *T. gondii* antigens from surviving mice. Dotted line indicates maximum limit of detection. **(G)** Brightfield images of vacuoles from WT, *Δgra72*, GRA72-complement, *Δgra17*, and GRA17-complement parasites. **(H)** Quantification of expanded vacuole occurrence across strains at 12, 24, and 30 h post infection. Mean ± SD indicated (*n* = 3, unpaired two-tailed *t*-test). **(I)** Representative images of CDCF vacuole-loading phenotypes from intracellular *Δgra17* parasites, incubated for 30 min with the dye. **(J)** CDCF vacuole-loading measured at 30 h post infection, incubating for 10 or 30 min with the dye. Mean ± SD indicated (chi-square test). **(K)** Immunofluorescence images showing the localization of GRA7 in WT and *Δgra72* vacuoles. **(L)** Immunofluorescence images showing cMyc staining of fibroblasts infected with WT, *Δgra72*, or GRA72-complement parasites. **(M)** Electron micrographs of fibroblasts infected with WT or *Δgra72* parasites 8 or 30 h post infection. Arrow indicates presence of host MLB. **(N)** Magnified image of host MLB in *Δgra72*-infected cell. **(O–P)** Quantification of the frequency **(O)** and area **(P)** of host MLBs in WT and *Δgra72* infected cells. Mean indicated (*n* = 3, unpaired two-tailed *t*-test).

We generated a knockout of *GRA72* to study its function (*Δgra72*; **Fig. 4C, S6B**). Despite forming normal plaques in cell culture, *Δgra72* parasites were avirulent in mice (**Fig. 4D–F**). While the PV membrane (PVM) is normally wrapped tightly around dividing parasites, *Δgra72* parasites displayed expanded vacuoles with large gaps between the parasites and the PVM (**Fig. 4G**). Complementation with a *GRA72-TY* construct rescued vacuole morphology and virulence in mice (**Fig. 4E–G, S6C**). Expanded vacuoles have been previously described in parasites lacking *GRA17*, which displayed a similar pattern to *GRA72* in our screen and has been shown to be required for mouse infection^69–71^ (**Fig. 4A**). We found that expanded vacuoles appeared earlier and in greater numbers in *Δgra72* parasites compared to *Δgra17* parasites (**Fig. 4G–H**).

Prior work used a low molecular-weight fluorescent dye (CDCF) to characterize GRA17 as a transporter of small molecules into the vacuole^69^. *Δgra72* vacuoles displayed a delay in dye uptake, consistent with decreased permeability to small molecules (**Fig. 4I–J**). While expanded vacuoles from both *Δgra72* and *Δgra17* parasites were impermeable to the dye during the first 10 min of incubation, expanded *Δgra72* vacuoles appeared permeable within 30 min. By contrast, *Δgra17* vacuoles showed minimal dye uptake with 30 min of incubation. Given their partial permeability and the higher frequency of expanded vacuoles in *Δgra72* parasites, GRA72 may have a broader impact on vacuole maintenance, rather than functioning as a transporter. Indeed, while GRA17 was first identified for its homology to the well-characterized *Plasmodium* transporter PfEXP2^69^, GRA72 bears no homology to known transporters.

We examined whether loss of GRA72 might impact secretion of effectors or their translocation out of the parasite vacuole. GRA7 and MAG1, two effectors that guard against antiparasitic host immune defenses^68, 72, 73^, were both secreted into the PV in the absence of GRA72 (**Fig. 4K, S6D**). Efficient effector secretion can also be assessed by its effect on host pathways. GRA16 is a well-characterized dense granule effector that induces nuclear localization of host c-Myc to suppress apoptosis of infected host cells^74^. *Δgra72* parasites were able to induce c-Myc nuclear staining, showing that *Δgra72* parasites secrete effectors normally (**Fig. 4L**).

We next used transmission electron microscopy (TEM) to examine the ultrastructural integrity of *Δgra72* vacuoles. The functions of the PV are supported by a series of tubular membranous structures called the intravacuolar network (IVN)^75, 76^. Despite their expanded morphology, *Δgra72* vacuoles had an intact IVN (**Fig. S7A**). However, *Δgra72* vacuoles displayed loose membranous structures peeling away from the inner wall of the PVM, suggesting a defect in PVM maintenance (**Fig. S7B**). *Toxoplasma* vacuoles also recruit host organelles to their surface^77–82^. *Δgra72* vacuoles displayed the typical association with host mitochondria, ER, and Golgi (**Fig. S7C**). However, we observed the formation of unexpected multilamellar bodies (MLBs) in the cytosol of *Δgra72*-infected host cells (**Fig. 4M–N**). To our knowledge, MLBs have not been previously reported as a consequence of infection, but are reminiscent of structures formed during autophagy^83, 84^. MLBs were more frequent and larger compared to any of the related structures found in WT-infected cells (**Fig. 4O–P**). *T. gondii* can trigger host autophagy to sequester host lipids through the recruitment of vesicles to the PV^85–87^. The buildup of MLBs and abnormal PVMs may therefore indicate an inability of these parasites to efficiently absorb host vesicles. Consistent with this model, we occasionally observed host organelles trapped within MLBs (**Fig. S7D**).

### GTP cyclohydrolase I is a druggable enzyme necessary during mouse infection

*T. gondii* depends on a broad and incompletely defined range of host metabolites for survival^30, 88, 89^. In surveying annotated metabolic genes, our screens revealed apparent differences in the pathways required to survive in mice compared to those required in cell culture. In addition to *PDX1* and *PDX2*, other enzymes required during mouse infection included those that participate in the metabolism of lipids (*TGGT1_242380* and TGGT1_257510)^90, 91^, pantothenate (*TGGT1_318600*)^92^, and glucose (*TGGT1_209960*)^93^ (**Fig. 5A**).

**Figure 5.**
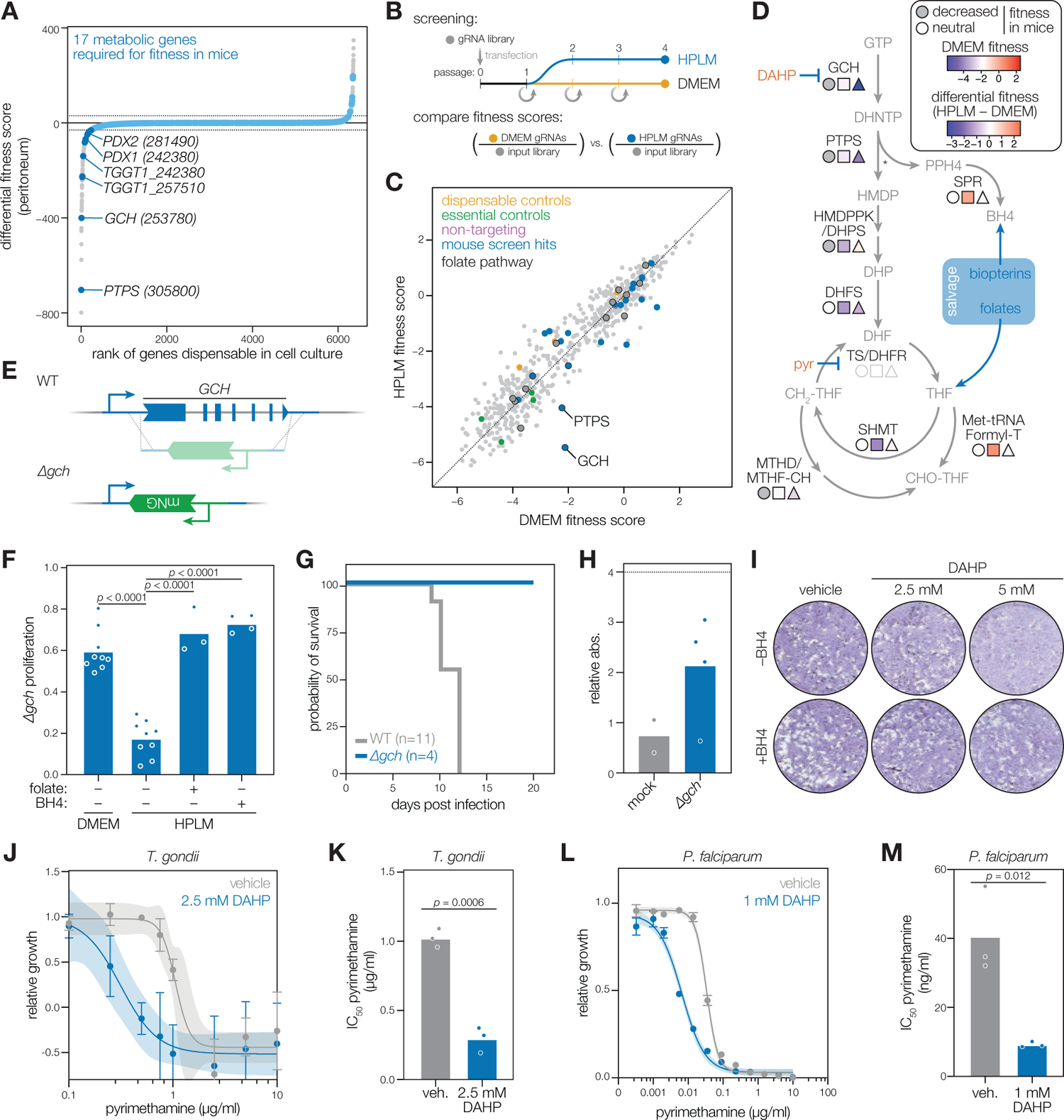
GTP cyclohydrolase I is a druggable enzyme necessary during mouse infection. **(A)** Rank-ordered plot of differential fitness for metabolic genes among all genes dispensable in cell culture. **(B)** Diagram of targeted metabolic screen in DMEM and HPLM. **(C)** Scatter plot of fitness scores of a targeted metabolic screen in DMEM and HPLM. **(D)** Diagram of folate and biopterin metabolism in *Toxoplasma.* Closed circles indicate a gene that scored with decreased fitness in mice. The shades of pink square and blue triangles indicate fitness score in DMEM and differential score (HPLM fitness - DMEM fitness) respectively. **(E)** Schematic of *GCH* knockout. **(F)** Relative fitness of *Δgch* parasites grown in DMEM, HPLM, or HPLM with folate and BH4 supplementation (unpaired two-tailed *t*-test). **(G)** Survival of CD-1 mice after infection with 10,000 WT or *Δgch* parasites. Number of mice in each group (n) is indicated. **(H)** Serum response to *T. gondii* antigens from surviving mice. Dotted line indicates maximum limit of detection. **(I)** Plaque assay of parasites grown in HPLM in the presence of DAHP and exogenous BH4 as indicated. **(J)** Dose response curve of *T. gondii* parasites grown in varying concentrations of pyrimethamine with and without 2.5mM DAHP co-treatment. Mean ± SD indicated. **(K)** IC_50_ value of pyrimethamine for *T. gondii* parasites with and without DAHP co-treatment (*n* = 3, unpaired two-tailed *t*-test). **(L)** Dose response curve of *P. falciparum* parasites grown in varying concentrations of pyrimethamine with and without 1mM DAHP co-treatment. Mean ± SD indicated. **(M)** IC_50_ value of pyrimethamine for *P. falciparum* parasites with and without DAHP co-treatment (*n* = 3, unpaired two-tailed *t*-test).

The dependency of parasites on genes like *PDX1* and *PDX2* during mouse infection highlights how nutrient concentrations in standard culture media can mask metabolic vulnerabilities in the parasite. Some differences between culture and animal environments can be reconciled by formulating media, like Human Plasma-Like Medium (HPLM), to mimic physiological nutrient concentrations^94^. To identify metabolic pathways in the parasite that respond to physiological media formulations, we conducted a metabolism-targeted CRISPR screen^30^ comparing standard culture conditions (DMEM) or HPLM (**Fig. 5B**). We found the majority of metabolic genes exhibited similar fitness scores in DMEM and HPLM, demonstrating that the metabolic challenges experienced by parasites during animal infection can not be easily captured in cell culture (**Fig. 5C, Table S9**). However, two candidates, GTP cyclohydrolase I (*GCH; TGGT1_253780*) and 6-pyruvoyl-tetrahydropterin synthase (*PTPS*; *TGGT1_305800*), were highly fitness-conferring in both HPLM media and during mouse infection. Both GCH and PTPS function in the metabolic pathways that produce tetrahydrobiopterin (BH4) and tetrahydrofolate (THF; **Fig. 5D**), two essential cofactors required for nucleic acid and amino acid metabolism^95, 96^. *GCH* is responsible for the conversion of GTP to 7,8-dihydroneopterin triphosphate. Biochemical analysis of the *P. falciparum* ortholog, suggests PTPS converts the product of GCH to either 6-pyruvoyl-5,6,7,8-tetrahydropterin (PPH4) or 6-hydroxymethyl-7,8-dihydropterin (HMDP), contributing to the production of both BH4 and THF^97^. While parasites and vertebrates share the ability to synthesize BH4, vertebrates depend on dietary folate to generate THF^31, 98–100^. By contrast, parasites can generate THF from both *de novo* biosynthesis and scavenging of biopterins and folates from the host cytosol (**Fig. 5D**)^31, 98–100^.

Folate is present at lower concentrations in HPLM compared to DMEM, potentially explaining the dispensability of parasite GCH and PTPS under standard growth conditions^94^. Under folate-limited conditions, our screen suggests that parasites rely more heavily on THF and BH4 biosynthesis. To test this hypothesis, we generated a *GCH* knockout (*Δgch*; **Fig. 5E, S8A**). As predicted by our screen, *Δgch* parasites showed a fitness-defect when cultured in HPLM. This defect could be rescued with the addition of exogenous folate or BH4 to the media (**Fig. 5F**). THF and BH4 are often described as endpoints of two biosynthetic pathways that branch after *GCH*. However, rescue of *Δgch* parasites by exogenously supplying folate or BH4 alone indicates that there is sufficient interconversion between these metabolites to sustain parasite replication. As vertebrate hosts are unable to synthesize folate themselves^99^, the interconversion of BH4 and folate must be completed within the parasite.

The reliance of apicomplexan parasites on folate has long been appreciated. Indeed, our screens detected several additional enzymes involved in folate metabolism as fitness-conferring during mouse infection, including dihydropteroate synthase (*DHPS, TGGT1_259550*) and tetrahydrofolate dehydrogenase (*MTHF; TGGT1_257625*). Current treatments for apicomplexan infections like toxoplasmosis and malaria rely on inhibition of folate metabolism through pyrimethamine and sulfadiazine, inhibitors of dihydrofolate reductase (DHFR) and DHPS, respectively^101^. DHFR functions downstream of the biosynthesis and salvage pathways that lead to THF^102^. DHPS is thought to function in folate biosynthesis^102^; however, the proposed interconversion of BH4 and folate implicates DHPS in a non-canonical salvage pathway. By contrast, GCH lies entirely upstream of folate and BH4 biosynthesis, without impacting salvage. Our results therefore indicate that, under physiological conditions, folate salvage alone is insufficient to support parasite growth. Indeed, *Δgch* parasites failed to cause infection in mice, despite intact salvage pathways (**Fig. 5G–H**). Chemical inhibition of *GCH* could therefore have antiparasitic effects.

2,4-Diamino-6-hydroxypyrimidine (DAHP), has been used to inhibit mammalian GCH for pain management, cancer, and cardiovascular disease^103–108^. A single report from 1964 documents the use of DAHP against the unrelated protozoan *Crithidia fasciculata*^109^, so we considered whether DAHP could be used against apicomplexans. Despite poor solubility and a high turn-over rate^106^, DAHP slowed parasite growth at a high dose, which appeared to be on-target based on rescue with exogenous BH4 (**Fig. 5I**). While we cannot rule out that antiparasitic effects of DAHP result from the inhibition of host GCH^110, 111^, the concentration of DAHP used did not impact proliferation of mouse fibroblasts (**Fig. S8B**). These results suggest there is potential for the development of GCH inhibitors as antiparasitic agents.

Clinical isolates of pyrimethamine-resistant *P. falciparum* were shown to have amplifications of the *GCH* locus, suggesting that increased flux through the folate biosynthesis pathway may facilitate pyrimethamine resistance^96, 112^. We therefore examined the efficacy of using DAHP in combination with pyrimethamine. DAHP and pyrimethamine were highly synergistic, leading to a 70% reduction in the pyrimethamine IC_50_ when cultures were supplemented with a sublethal dose of DAHP (**Fig. 5J–K**). Synergy between the DAHP and pyrimethamine was also found in *P. falciparum*, as co-treatment resulted in a 77% reduction in the IC_50_ of pyrimethamine (**Fig. 5L– M**). By contrast, DAHP showed no synergy with atovaquone, an inhibitor of the cytochrome bc1 complex unrelated to folate metabolism^113^ (**Fig. S8C–D**). While further optimization is needed to improve the antiparasitic efficacy of DAHP as a single agent, the synergy of GCH inhibition with pyrimethamine may help prevent the emergence of resistance and reduce patient toxicity from standard pyrimethamine treatments.

## DISCUSSION

In this study, we present the first genome-wide analysis of an apicomplexan during infection of an animal host. We identified over 300 genes that impact fitness during mouse infection, most of which remain uncharacterized. Demonstrating the power of our screen to identify new axes of host-parasite interaction, we functionally characterize three novel candidates that highlight features of host cell tropism, remodeling of the replicative niche, and metabolic rewiring.

Our screen employed barcoded gRNAs to track parasite populations during mouse infection. Barcodes have been used to map population structure in studies of viral^114–117^, bacterial^118–125^, and parasitic infections^126, 127^, revealing host tissues that are permissive or restrictive to pathogen colonization and growth. A recent report in *T. gondii* used barcoded parasites to demonstrate that a limited number of clones reach the brain, with considerable stochasticity in clonal makeup across individual mice^126^—a pattern also documented in our data. Combining barcoding with genome-wide knockouts, our study expands on this approach by tracing population structures across all single-gene parasite mutants. In addition to tracking mutant dissemination patterns, we use these barcodes to denoise our screens by identifying bottlenecks and jackpotting artifacts through a broadly applicable analytical framework. While barcodes have been used to improve screening efficiency in other contexts, previous methods rely on consistent detection of UMI-gRNA pairs^18–21^, which can prove difficult in sparse datasets. By relying instead on population-level metrics, our method facilitates exclusion of unreliable gRNA scores even in any highly bottlenecked screening contexts.

The combination of barcoded gRNAs and improved screening methods significantly increased the accuracy and efficiency of *T. gondii* screening in mice. Comparing our results with previous small-scale screens, we found five of the 24 genes identified by Sangaré et. al. and three of the eight identified by Young et al.^14, 15^. Differences in the way the screens were conducted and analyzed likely contribute to these discrepancies. We observed seven additional hits from the prior screens (five from Sangaré et. al. and two from Young et al.) were progressively lost in cell culture and therefore excluded from our set of candidates. This highlights the importance of time-matched comparisons between cell culture and mouse samples to avoid false positives. The improved dynamic range and reproducibility of our screen—approximately three times greater than that of previous screens—helped us better distinguish between biological effects and statistical noise. Additionally, Young et. al. conducted their study using a less virulent type II strain (PRU) in inbred C57Bl/6J mice, as opposed to a type I strain in outbred CD-1 mice. We can therefore attribute some discrepancies between our screens to the host and parasite backgrounds; e.g., ROP18 (*TGGT1_205250*) confers protection from host immunity in type II strains, but its loss in type I strains only marginally delays mouse morbidity^27^. Consequently, *ROP18* displayed a neutral fitness score in our screen. Overlap between screens provides a strong argument for standardizing screening methods, while screening details and quantitative metrics can help determine which results should guide further experimental validation.

In exploring our candidate set, we characterized three genes with decreased fitness in mice that span diverse aspects of the host-parasite interface. We first identified RASP1 as a mediator of repeated invasion, a process that remains poorly understood in *Toxoplasma* parasites. RASP1 is among a number of genes identified in our screen that function to support the basic mechanics of the parasite lytic cycle, yet display only neutral or moderate phenotype scores in cell culture. Other factors implicated in invasion (SPATR^33^, ROM4^45^, TgNd6, and TgNdP2^128^) and egress (PLP1^34^) were also identified by our screen, demonstrating that even fundamental steps in the lytic cycle are pressure-tested during whole-organism infections in ways that are incompletely captured in cell culture. The macrophage-specific defect of *Δrasp1* parasites suggests that broad host cell tropism may require multiple rounds of rhoptry discharge. Permissiveness to parasite invasion may arise from differences in host membrane composition, such as cholesterol content^129^. Additionally, secreted rhoptry proteins inactivate host antiparasitic pathways. Host cells with elevated innate immunity, such as macrophages, may require multiple rounds of rhoptry protein secretion for parasites to adequately establish a replicative niche^130^.

Rhoptry proteins must be translocated across three lipid bilayers: the bounding membrane of the rhoptry, and the plasma membranes of the parasite and host cell. The mechanics of rhoptry-content secretion and the turnover of discharged rhoptries remain under active investigation. Cryo-electron tomography studies have recently clarified the ultrastructure of the invasion machinery by identifying apical vesicular structures that connect docked rhoptries to the parasite plasma membrane^65^. Fusion of rhoptries with the apical vesicle appears to depend on the RASP1 paralog, RASP2^58^. *T. gondii* tachyzoites contain multiple rhoptries and apical vesicles^65^, so RASP1 may facilitate fusion of undocked rhoptries or mediate turnover of discharged rhoptries. To our knowledge, RASP1 represents the first factor implicated in repeated invasion attempts. Future cryo-electron tomography studies of *Δrasp1* parasites may provide further details on the reloading of the invasion machinery.

We also characterized a dense granule protein, GRA72, involved in PV maintenance and virulence in mice. Several *T. gondii* factors aid in remodeling host cell organelles and vesicular trafficking. MAF1, GRA3, GRA14, and GRA64 all play distinct roles in recruiting mitochondria, Golgi material, and the ESCRT machinery to the PV^79, 80, 131, 132^; however, in contrast to GRA72, none of these factors appear to contribute to parasite fitness in cell culture nor in mice during the acute phase of infection. Loss of GRA72 therefore seems to interfere with a distinct host-remodeling process that is crucial to parasite viability in mice. Large MLBs formed in the cytosol of *Δgra72*-infected host cells. MLBs have been previously associated with autophagy induced by ER stress. For example, neuroblastoma cells treated with the ER stressors tunicamycin or thapsigargin develop MLBs within hours, while starvation in these cells only induced canonical autophagosomes^84^. *Toxoplasma* infection induces host ER stress, suggesting a possible origin for the MLBs found in *Δgra72*-infected cells^133–135^. Our discovery of GRA72 sheds light on the challenge of maintaining the function of a growing PV, and how this process may be further strained in the organismal context.

We also characterized specific metabolic challenges encountered by parasites in mice. A media formulation that mimics nutrient availability in human plasma (HPLM) could recapitulate only a few of the metabolic dependencies identified in mice. This indicates that HPLM provides an incomplete representation of the nutrient environments experienced by parasites during mouse infection. These discrepancies may arise from cell-type specific metabolism^136, 137^, or alterations to nutrient levels as a result of inflammation^138–141^. Investigation of other metabolic dependencies identified by our screen will help define the nutrient environments experienced by parasites during infection.

Our characterization of *GCH* demonstrates the potential to identify new druggable parasite vulnerabilities that arise during animal infection. We found that a previously characterized inhibitor of mammalian GCH (DAHP), exhibits an antiparasitic effect. The pharmacokinetics of DAHP limit its use to proof-of-concept studies, requiring more potent inhibitors be developed for clinical use. Substantial sequence divergence between parasite and human GCH suggests it may be possible to develop parasite-specific inhibitors. We show that GCH inhibitors may be clinically appealing for their ability to synergize with established antifolate drugs, such as pyrimethamine. While these drugs are considered frontline treatments for apicomplexan infections, extended administration can result in patient toxicity^142^. Additionally, resistance is emerging in *Plasmodium* spp.—in some cases through *GCH* copy number amplifications^96, 112^.

Our study of GCH also revealed the previously unappreciated interconversion of BH4 and folate in parasites, which likely contributes to DAHP synergy with antifolate drugs. Mammals are incapable of folate biosynthesis, yet retain GCH-mediated production of BH4, suggesting parasites may scavenge host BH4 to synthesize folate under limiting conditions. A parasite-specific GCH inhibitor may therefore prove effective on its own or in combination with established antifolates.

In addition to identifying genes that contribute to parasite fitness in mice, our screen revealed 111 genes that increased fitness when disrupted. While initially unexpected, several recent reports have identified factors that enhance pathogen fitness under specific conditions when deleted. Screens targeting *Leishmania mexicana* kinases and *Legionella pneumophila* effectors identified analogous groups of genes that improved pathogen fitness during mouse infection when disrupted^143, 144^. Enhanced fitness may arise through various mechanisms. Pathogen effectors may be recognized by host immunity. For example, the polymorphic secreted protein GRA15 recruited host ubiquitin ligases to the PV in certain *T. gondii* strains, and its loss conferred a relative fitness advantage over WT parasites during infection of interferon-stimulated host cells^145^. Similarly, recognition of flagellar proteins mediates host restriction of *L. pneumophila* by inflammasome activation, leading to increased fitness of flagellin-deficient mutants^144, 146–148^. Loss of certain genes may only be feasible in a pooled setting, where mutants still benefit from the effect of wild-type organisms. Expression of a type III secretion system 1 (ttss-1) allows *Salmonella typhimurium* to outcompete commensal bacteria by inducing host inflammation^149^. *Δttss-1* mutants are avirulent alone, but can outcompete wildtype *S. typhimurium* in a mixed population by avoiding the fitness cost of expressing the secretion system, while reaping the benefits of its expression. Increased fitness may also be driven by pathogen-intrinsic processes, such as preventing transitions to slower-replicating life stages. For example, loss of the ApiAP2-G transcription factor in *P. berghei* prevents differentiation to the non-proliferative sexual stages and increases parasitemia through continued asexual replication^150^. Finally, a portion of the increased fitness candidates may arise from defects in cell culture that are rescued in the animal host, as is the case with candidates involved in oxidative-stress responses. In all scenarios, functional characterization of these candidates will likely point to new context-specific demands on parasite biology and host modulation.

The improved screening approaches we have described will help unravel host-parasite interactions in diverse host contexts. For example, susceptibility to *T. gondii* infection varies across mouse genotypes. Much of this susceptibility is attributed to polymorphic immune-regulated GTPases that are coevolving with parasite effectors^151, 152^; however, screens in different mouse backgrounds may uncover additional genotype-specific facets of host-parasite interaction. Screens in mice depleted for certain immune lineages or lacking particular cytokines may also reveal parasite effectors that guard against specific host defenses^153^. While we have focused on the earliest stages of infection, similar methods could be deployed to understand other life-cycle stages or alternative infection routes. Previous screens examining the chronic phase of *T. gondii* infection have relied on experimental induction of the chronic stages in cell culture^154^. While we found the brain to be highly bottlenecked in genome-wide screens, smaller libraries would enable dissection of the chronic stage in a tissue where it naturally develops. Further, the broad tropism of *T. gondii* across warm-blooded animals may have driven the evolution of parasite genes required in specific host species^6^; genes identified as dispensable during mouse infection could still prove to be fitness-conferring in other animals.

Our work represents an unprecedented resource, mapping the fitness landscape of all single-gene mutants in a eukaryotic pathogen during the infection of its natural host. Apicomplexan parasites remain understudied organisms despite their relevance as human and animal pathogens. While host species and life cycles of apicomplexans vary, many of the genes that modulate *T. gondii* fitness in mice are broadly conserved across the phylum and likely participate in general adaptations to host pressures. *T. gondii* has often served as a representative of the apicomplexan phylum for its ease of culturing and genetic manipulation—our work adds genome-wide functional genomics during animal infection as a new tool to investigate the adaptations that mediate apicomplexan parasitism.

## MATERIALS & METHODS

### gRNA library design

Nine-gRNA and three-gRNA genome-wide libraries were designed by a proprietary service provider according to current guidelines for gRNA efficiency^22^. gRNAs for the 17 sub genome-wide libraries were subsetted from the three-gRNA genome-wide library. Sublibraries 1–5 were designed around specific target gene sets of interest (**Table S1**). Key words for rhoptry, dense granule, and microneme were used to generate gene lists from ToxoDB.org. Additional genes of interest were identified from an unbiased proteomics-based localization dataset^52^. Proteins with predicted signal peptides were also identified from ToxoDB.org. The remainder of the genome was randomly divided between the remaining 12 sublibraries. In an effort to promote similar screening dynamics across libraries, small adjustments were made in gene sets among these 12 sublibraries to ensure that each had a similar mean and standard deviation of gene fitness scores as informed by our three-gRNA genome-wide screen. 199 genes had been left out of the genome-wide gRNA libraries due to poor genome annotation or difficulty in identifying gene-specific gRNAs. gRNAs against these genes were manually generated and confirmed for gene specificity at a cut off of at least 4 mismatches in potential off-target genes. 31 gRNAs remained in our library with multiple targets, often targeting duplicated genes or poorly mapped genes present on contigs (**Table S1**). gRNAs were checked for absence of an AseI cut site, as this would interfere with library construction, and swapped for an alternative gRNA sequence when needed. Each library also included 30 nontargeting gRNAs and control gRNAs targeting genes known to be fitness-conferring in cell culture or mice. Control genes known to be fitness-conferring in cell culture included *TGGT1_293690*, *TGGT1_253040*, *TGGT1_209030*, *TGGT1_313380*, *TGGT1_228750*, *TGGT1_243250*, *TGGT1_235470*, and *TGGT1_256920*. Control genes known to be fitness-conferring in mice included *TGGT1_243730* and *TGGT1_316250*.

### gRNA library construction

gRNA libraries were synthesized in pools by Agilent with barcoded ends allowing for amplification of specific sublibraries. First, primers unique to each sublibrary barcode were used to amplify a single sublibrary at a time from the pool using iProof polymerase (**Table S10**). Amplified libraries were then gel extracted and purified with the Zymoclean Gel DNA Recovery Kit. Amplified libraries were then diluted to 2 ng/µl and PCR amplified with iProof with internal primers matching the homology region for subsequent gibson assembly into the gRNA vector backbone. PCR products were gel extracted and purified.

Separately, gRNA vector backbones were digested with BsaI-HF V2 overnight and purified by PCR cleanup. Purified digested vector was digested a second time with BsaI-HF V2 for 2 h. In the last hour of incubation, CIP was added to decrease the rate of self-ligation of the vector. The double-digested vector was again purified by gel extraction. Mouse screen sublibraries were cloned into UMI-barcoded gRNA vector backbone. Methods for construction of the UMI-barcoded gRNA vector backbone are described below.

Gibson assembly of a gRNA library into the gRNA backbone was completed with NEB HiFi 2x mastermix with a 1:5 molar ratio of vector to insert in 10 µl total volume. After assembly reactions were completed, products were dialyzed against water for 20 min on 0.025 µm MCE filter membranes.

Prior to bacterial transformation, large agar plates were poured and cooled containing 100 µg/ml carbenicillin. Assembled libraries were electroporated into MegaX DH10B T1R Electrocomp Cells (Invitrogen C6400-03) with 10 µl divided over 5 transformations of 20 µl of cells each (electroporated at 2000 V, 200 ohms, 25 µF). Transformed cells were incubated in SOC at 37°C for 90 min, then spread over 5 large agar plates. After 18 h of growth, bacterial colonies were scraped from agar plates, pelleted, and frozen at −20°C. Plasmids were later harvested from bacteria pellet by MN Maxi Kit.

*UMI-barcoded gRNA vector construction:* Mouse screen libraries were gibson assembled into digested UMI-barcoded gRNA backbones. This barcoded backbone was generated by amplifying a ∼100 bp fragment at the end of the gRNA scaffold with a primer containing 8 internal sites of randomized base pairs (**Table S10**). This PCR product was gel extracted and purified with the Zymoclean Gel DNA Recovery Kit. Separately, 5 µg of the parental plasmid was digested with NsiI and NdeI, which cut at sites with homology to the gRNA amplicon. The digested vector was gel purified, and digested a second time with NsiI and NdeI for 2 h, followed by CIP treatment. NEB 2x HiFi master mix was used to assemble the short fragment containing UMIs to the digested vector. This product was then transformed into MegaX DH10B T1R Electrocomp Cells by electroporation as described above.

### Nine-gRNA and three-gRNA genome-wide CRISPR screens

Genome-wide screens using the nine-gRNA and three-gRNA libraries were conducted as previously described with modifications^155^. Briefly, gRNA libraries were linearized with AseI prior to transfection. The digest library was then transfected into Cas9-expressing parasites^156^ and infected across 10 x 15 cm dishes of confluent HFF cells. One day after transfection, plates were washed with PBS and replenished with media containing 10 µg/ml DNaseI and 3 µg/ml pyrimethamine. Every two days from the transfection, parasites were scraped, mechanically lysed, and filtered through 0.5 µM filters. At passage 4, 1 x 10^8^ parasites from each sample were pelleted and frozen at −80°C. Genomic DNA was harvested from these samples using the Qiagen DNeasy Blood & Tissue kit. gRNAs were amplified from genomic DNA as well as the transfection input using Q5 polymerase master mix. Amplified products were gel extracted and purified with the Zymoclean Gel DNA Recovery Kit. gRNA abundances were quantified by Illumina sequencing (**Table S2**). gRNAs of the lowest 1% abundance of the input library were filtered from downstream analyses. Gene scores were calculated as the mean log_2_(fold change) of gRNA abundance at the indicated sample relative to the transfection input.

### Mouse screens

Mouse screens were conducted similarly to the nine-gRNA and three-gRNA genome-wide CRISPR screens, with modifications. Briefly, gRNA libraries were linearized with AseI prior to transfection. Two sub genome-wide libraries were pooled in a single screen (**Table S1**). The digested pooled sublibraries were transfected into Cas9-expressing parasites^156^ and infected onto 8 x 15 cm plates of confluent HFFs. One day after transfection, plates were washed with PBS and replenished with media containing 10 µg/ml DNaseI and 1.5 µg/ml pyrimethamine. Every two days until the end of the screen, parasites were scraped, mechanically lysed, and infected onto 8 new HFF monolayers. Samples of 1 x 10^8^ parasites were harvested at each passage after passage 4.

At passage 4, part of the population was pelleted and washed in PBS, then pelleted again and resuspended to a concentration of 1 x 10^8^ parasites per ml. 1 x 10^7^ parasites were injected into 10 female CD-1 mice of 8-10 weeks of age. Six days after infection, mice were euthanized. Parasites were harvested from the peritoneum by peritoneal wash with 5 ml of 10% IFS in PBS. Solid tissues (liver, spleen, lung, heart, and brain), were dissected out and homogenized in a 12-well plate with the back of a plunger syringe. Homogenized tissues were then pushed through 70 µm filters and mechanically lysed through 27 Ga needles. Peritoneum, spleen, lung, and heart samples were then plated onto 15 cm of confluent HFFs, while samples from liver and brain were divided across two 15 cm plates each. Tissue harvest plates were maintained in DMEM supplemented with 10% IFS, 2 mM glutamine, 10 µg/ml gentamicin, and 1x Penstrep. In our pilot screen, prior to plating, half of the sample was reserved for direct genomic DNA harvest without an outgrowth step. Infected dishes were monitored for lysis. Peritoneal samples tended to lyse out 2 days after infection, while liver, spleen, and lung samples would lyse 3–4 days after infection, and heart and brain samples would lyse 5–6 days after infection. Lysed plates were scraped, passed through a 27 Ga syringe, then pelleted and stored in −80°C. Genomic DNA was harvested from lysed plates infected from peritoneum, liver, spleen, lung, and heart samples with the Qiagen DNeasy Blood & Tissue kit. Genomic DNA was harvested from lysed plates infected with brain samples with the Zymo Quick-DNA Midiprep Plus Kit to account for the greater amount of cell debris in those samples. gRNAs were amplified from genomic samples with

NEBNext Ultra II Q5 master mix. Amplified products were gel extracted and purified with the Zymoclean Gel DNA Recovery Kit. gRNA abundances (read 1) and UMI barcodes (read 2) were quantified by Illumina sequencing and subsequently processed by the analysis described below.

#### Barcoded CRISPR screen analysis

All subscreens were analyzed separately before pooling results at the final stage of analysis (**Table S5**).

#### Filtering for read quality

Read 2 reads were first filtered for length by removing reads with UMIs shorter than 8 bps. The remaining reads were then filtered by a q-score of at least 28. Remaining read 2 UMI reads were then paired with their read 1 gRNA counterparts.

#### Quantifying gRNA abundance

Reads from each sample were then mapped to the gRNA library to quantify relative gRNA abundances. For each gRNA from each sample, the relative contribution of a barcode to the gRNA abundance was recorded.

#### Generating initial gene fitness scores

gRNA abundances were normalized as the fraction of total reads for each sample. The gRNAs in the lowest 1% of abundance of the input library were removed from subsequent analysis. Log_2_(fold changes) were calculated for each gRNA from each sample relative to the input library. Gene fitness scores were calculated as the average gRNA score for all gRNAs against that gene.

#### Removing outliers from high variance genes

To filter out gRNAs that show consistently poor on-target or off-target activity, we identified genes with the highest variance between their gRNA scores. As disruption of a gene may only show a phenotype in mice but not in cell culture, we applied variance filtering to high-coverage samples from multiple stages of our screen (passage 8, mouse peritoneum, liver, and lung). Since mice are independently infected, we had an additional requirement that a gene must be among the top 5% in variance for at least *n*-1 mice for the subscreen (*n* = total number of mice used in that screen). Once nominated as a gene with high variance, the outlier gRNA in the set was removed by comparing gRNA scores generated from pooled mouse samples. In this analysis, we summed normalized raw reads across all mice for a given tissue, and computed gRNA log_2_(fold changes). We then identified the outlier gRNA among the set of three, and excluded it from downstream analysis.

#### Incorporating barcode diversity metrics

We next set out to identify instances of bottlenecking and jackpotting in our screen by analyzing the distribution of barcodes for each gRNA in each sample. We began by filtering out barcodes that appeared only once for a well-represented gRNA population, as these are likely to arise from sequencing errors or template switching^157^ during the gRNA amplification from genomic DNA. For each gRNA in each sample, we calculated species richness and Shannon evenness. Species richness is simply the total number of unique barcodes identified for that gRNA in that sample. Shannon evenness was calculated as follows:

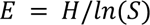

Where *E* is Shannon evenness, *S* is species richness, and *H* is Shannon diversity as defined below:

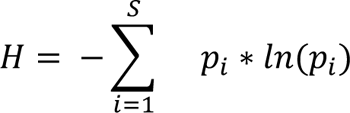

#### ML-generation of evenness outlier thresholds

We observed that the distribution of evenness varied with respect to gRNA abundance. We therefore used dynamic thresholding to identify gRNAs with outlier evenness scores relative to gRNAs with similar log_2_(fold changes). We used a machine learning Bayesian model to set thresholds as 2 standard deviations below the predicted population modeled by all samples for each tissue within a subscreen. Outlier gRNAs within each sample were then excluded from downstream analysis.

### Gene fitness scores and differential fitness

Having removed variance and evenness outliers from our data, we next generated final gRNA log_2_(fold changes) relative to the input library. Gene fitness scores were calculated as the mean gRNA score for all gRNAs targeting a gene. In cases where a gene was screened in multiple subscreens (as a control included in all libraries or a replicate screen), gene fitness scores were averaged. Heart and brain samples were particularly bottlenecked. We therefore pooled normalized reads from individual mice into one “metamouse” for each subscreen for heart and brain samples, and processed these combined reads as a single mouse sample.

The peritoneum was used as the reference tissue to calculate differential fitness between mice and cell culture. For each mouse from each subscreen, a linear regression was modeled for peritoneal gRNA scores against cell culture passage 8 scores. This model was used to generate a distance metric from the “predicted” peritoneal gRNA value from the observed peritoneal gRNA value. Distance metrics calculated for nontargeting gRNAs from each mouse in a subscreen were pooled to generate a null distribution of gRNA distances. Two-tailed *t*-tests were then used to compare the distribution of distances of gRNAs against a given gene across all mice in a subscreen relative to the null distribution.

### Tissue differential fitness

Differential fitness comparisons between tissues were calculated similarly to the reference comparison of peritoneum scores against cell culture. In these comparisons, linear regressions were computed for each mouse within a subscreen between the two tissues of interest (*x* = tissue_1 and *y* = tissue_2). This model was used to generate a distance metric from the “predicted” tissue_2 gRNA value from the observed tissue_2 gRNA value. Distance metrics calculated for nontargeting gRNAs from each mouse in a subscreen were pooled to generate a null distribution of gRNA distances. Two-tailed *t*-tests were then used to compare the distribution of distances of gRNAs against a given gene across all mice in a subscreen relative to the null distribution.

### Genetic distance, Jaccard index, and tissue dendrogram generation

Nontargeting gRNA barcodes were used to analyze barcode population dynamics between tissue sites. Barcodes were first filtered to those initially identified in the input library sample for each subscreen to filter barcodes that may have been introduced by sequencing artifacts. The fraction of UMIs shared between two tissue sites were calculated as the Jaccard index of barcode identity between two tissue sites for a nontargeting gRNA within a mouse^158^. Genetic distance between two tissues was calculated for each mouse, taking into account all nontargetting gRNAs and their barcodes as features of the population in a single metric^159^:

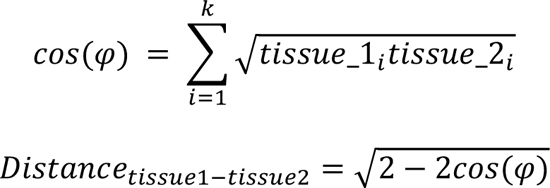

Where *tissue_1* and *tissue_2* are the two tissues being compared. k is the total number of unique UMI-gRNA pairs identified from either tissue. *tissue_1_i_* and *tissue_2_i_* refers the relative frequency of the particular UMI-gRNA pair (*i*) among the total nontargeting UMI-gRNA read counts. Dendrograms between tissues were generated for each mouse based on genetic distance using the neighbor joining method using the APE^160^ and phangorn R^161^ packages.

### Gene set enrichment analysis

Gene set enrichment analysis was conducted using published gene localization assignments^52^. The hypeR R package^162^ was used to run hypergeometric tests to identify significantly enriched categories among the 348 differential fitness candidates. Enrichment fold change was calculated as the ratio of the fraction of a category of interest present within the candidate list relative to that fraction in the background genome.

### Cell culture

Type I RH parasites (ATCC PRA-319) were grown in human foreskin fibroblasts (HFFs, ATCC SCRC-1041) and maintained in DMEM (GIBCO) supplemented with 2 mM glutamine, 10% inactivated fetal bovine serum (IFS), and 10 µg/ml gentamicin. C57Bl/6J mouse embryonic fibroblasts (B6 MEFs) and RAW 264.7 macrophages were maintained in DMEM (GIBCO) supplemented with 2 mM glutamine, 10% IFS, and 10 µg/ml gentamicin (Thermo Fisher Scientific). When indicated, Human Plasma-like Media (HPLM, GIBCO) was used as base media and supplemented with 2 mM glutamine, 10% IFS, and 10 µg/ml gentamicin.

The NF54 *P. falciparum* line was cultured and synchronized as described previously^163, 164^. Cultures were kept at 37°C in RPMI-1640 medium supplemented with 25 mM HEPES, 100 μM hypoxanthine, 0.5% Albumax II, 24 mM sodium bicarbonate, 25 μg/mL gentamicin (Life Technologies, Carlsbad, CA 11021-045) and gassed with 5% CO_2_, 1% O_2_, and 94% N_2_ mixture.

### Mice

All experiments were conducted in 8-10 week old female CD-1 (Charles River) mice. Mice were maintained in Whitehead Institute facilities accredited by the Association for Assessment and Accreditation of Laboratory Animal Care. Protocols were approved by the Institutional Animal Care and Use Committee at the Massachusetts Institute of Technology.

### Plaque assays

Intracellular parasites were mechanically-lysed and filtered. 500 parasites of the indicated strain were infected into a well of a 6-well plate in the indicated media. For DAHP plaque assays, wells were then treated with the indicated concentrations of DAHP or PBS vehicle. For BH4 rescue experiments, wells received 50 µM BH4 or the equivalent volume of DMSO vehicle. Plaques were allowed to form for 7 days prior to PBS wash and ethanol fixation. Wells were stained for 10 min with crystal violet (12.5 g crystal violet, 125 ml 100% ethanol, 500 ml 1% ammonium oxalate). Wells were then washed three times with water and allowed to dry overnight. Stained plaques were scanned on an Epson Perfection V800 Photo scanner. All plaque assays were conducted at least twice.

### Parasite transfection and strain construction

Existing RH Type I parasites were used as the parental strain for all strains generated in this study. All parasite transfections were completed in parasites resuspended in transfection solution containing 0.15 mM CaCl_2_, 2 mM ATP, 5 mM glutathione, 15–60 µg of DNA, and cytomix (10 mM KPO_4_, 120 mM KCl, 5 mM MgCl_2_, 25 mM HEPES, 2 mM EDTA, 2 mM ATP, and 5 mM glutathione) up to 400 µl. Parasites were transfected by electroporation with two 150 µs pulses at a 100 ms interval at 1700 V.

### Generating knockout parasites

gRNAs targeting the 5′ and 3′ ends of each gene were cloned into a Cas9-expressing vector containing a sgRNA scaffold. A pTubulin_mNeonGreen_3′*DHFR-*UTR expression construct was PCR amplified with overhangs homologous to the regions immediately surrounding the cut sites of these gRNAs. The two gRNAs and the mNeonGreen repair template were transfected into RH*Δku80Δhxprt* or RH*Δku80Δhxgprt::nanoLuc* parasites. At first lysis (48 h), transfectants were FACS sorted for mNeonGreen positive parasites. Parasites were then subcloned into 96 well plates. After 6–7 days of growth, wells containing single plaques were selected. Single plaques were expanded and PCR validated for successful knockout of the gene of interest (see **Table S10** for relevant primers for strain construction).

### Endogenous tagging of RASP1, GRA72, and MAG1

Genes were endogenously tagged using the previously described high-throughput (HiT) tagging system^165^. Cutting units targeting the 3′ coding sequence ends of RASP1, GRA72, and MAG1 were synthesized from IDT and cloned into HA or TY-epitope tagging HiT vector backbones. Prior to transfection, HiT vectors were linearized with BsaI-HF V2. Between 15–60 µg of digested plasmid was co-tranfected with a Cas9-expressing vector into RH*Δku80/Δhxgprt* parasites. One day post transfection, parasites were selected with 1 µM pyrimethamine or 25 µg/mL mycophenolic acid and 50 µg/mL xanthine pH-balanced with 0.3 N HCl. After approximately 1 week of selection, parasites were dilution plated into 96 well plates to obtain single parasite clonal populations. Clonal populations were then PCR verified for correct HiT vector integration and assessed by IFA for presence of the epitope tag.

### Generating complemented strains

Complementing plasmids for RASP1 and GRA72 were constructed with the endogenous UTR sequences, intron-less coding sequences, C-terminal TY epitope tags, and a pyrimethamine drug resistance cassette (**Table S10**). For transfection, complementing repair templates were PCR’d with overhangs bearing homology to an intergenic neutral locus in the parasite genome^61^. For transfection, *Δrasp1* or *Δgra72* parasites were transfected with their respective complementing templates along with a gRNA targeting the intergenic locus.

### Immunofluorescence Assays

Parasites were infected onto confluent HFF coverslips for the specified period of time. At time of fixation, coverslips were washed 3x with PBS, then fixed with a 4% formaldehyde solution at 25°C. After 20 min, the 4% formaldehyde was removed and the coverslips were washed 3x with PBS. Wells were then permeabilized with 1% triton for 8 minutes, then washed 3x with PBS. Wells were then blocked for 10 min with a 5%-NGS/5%-IFS PBS blocking solution. Primary antibodies were incubated on coverslips at the following dilutions for 30 min at 25°C. After 30 min incubation, wells were washed 3x with PBS, and then blocked for 10 min with 5%-NGS/5%-IFS, then incubated with the appropriate secondary antibody and hoechst for 30 min. After the incubation, wells were washed 3x with PBS, dipped 2x in dH_2_O, and mounted on slides with Prolong Diamond Antifade Mountant until the mounting media solidified. Coverslips were imaged on an Eclipse Ti microscope (Nikon). See **Table S10** for relevant antibodies and dilutions.

### Mouse competition infection assays

Competitions were conducted with green KO strains and red WT strains. Intracellular parasites were scraped and mechanically lysed through 27 Ga needles. Parasites were passed through a 5 µM filter to remove host cell debris prior to counting. 8 x 10^5^ parasites of each WT and KO strains were combined, and checked for an approximate 1:1 mix by flow cytometry. After confirmation of the starting ratios, parasites were infected onto 1 well of a 6 well plate, and spun at 290 x *g* for 5 min. The competition was passaged in culture for at least 2 passages prior to mouse infection, and measured by flow cytometry for ratios of KO and WT parasites. On the day of mouse injection, intracellular parasites were washed with PBS. Wells were scraped, mechanically lysed, and filtered. Ratios of KO:WT parasites were measured by flow cytometry and recorded as the input prior to mouse injection. Filtered parasite pools were diluted to 5 x 10^6^ parasites per 100 µl in PBS. 8–10 week-old CD-1 female mice were then injected IP with 100 µl of the parasite pool. Mice were sacrificed 6 days post infection, and parasites were harvested from the peritoneum and lung as described in the mouse screen. Harvested parasites were infected onto confluent HFF 15 cm plates, and allowed an outgrowth period of 1–2 days. At the end point, ratios of KO to WT parasites were measured by flow cytometry from the harvested parasites and compared to ratios of the parasite pool maintained in cell culture.

### Serum ELISA for anti-Toxoplasma antibodies

Blood was taken from infected mice 6–8 weeks post infection by tail bleed. Blood was set to coagulate at 25°C for 30 min, and the centrifuged at 2000 x *g* at 4°C for 15 min. The soluble serum fraction was then stored at −80°C until ELISA analysis. Soluble Toxoplasma antigen (STAg) was generated as previously described^166, 167^. On day 1, 96 well flat bottom plates were coated with 50 µl/well of 5 µg/ml STAg and incubated overnight at 4°C. On day 2, plates were washed 3x with ELISA wash buffer (0.5% Tween in PBS) and blocked for 2 h at room temperature with 10% IFS. Serum was then added to wells at 1:10, 1:50, 1:200, and 1:400 dilutions, and incubated overnight at 4°C. On day 3, wells were washed 3x with wash buffer, then incubated with anti-mouse FC antibody at 1:500 in blocking buffer for 2 h. Wells were then washed 3x with wash buffer, and mixed with 100 µl of TMB solution and incubated at room temperature for 2 min. 100 µl of stop solution (1M HCl) was then added per well. Wells were imaged for absorbance at 450 nm.

### Generation of CD-1 bone marrow-derived macrophages

Bone marrow macrophages were obtained from CD-1 female mice. Mice were euthanized, and femurs and tibias were dissected out. A 18 Ga needle was used to push out bone marrow with DMEM supplemented with 10% IFS. Bone marrow was resuspended in 5 ml of DMEM + 10% IFS. Cells were then pelleted at 290 x *g* at 4°C for 5 min. Cells pellets were then resuspended in 10 ml of macrophage media (DMEM supplemented with 10% FBS, 1% sodium pyruvate, 20 ng/ml mouse macrophage colony-stimulating factor) and kept on ice. Cells were counted, and 5 x 10^6^ cells were plated in 10 cm bacterial Petri dishes (non tissue-culture treated), and incubated at 37°C with 5% CO_2_. After 3 days, 4 ml of fresh macrophage media was added to each plate. Macrophages were harvested 6 days after initial plating. Plates were kept on ice for 20 min, and media was removed and replaced with 10 ml of cold PBS. Macrophages were gently scraped off of the plate, and pelleted at 4°C at 290 x *g* for 5 min. Pellets were then resuspended to the desired concentration for re-plating for experiments, or suspended to 1 x 10^6^ cells per ml in freezing media (Macrophage media supplemented with 10% DMSO) for freezing.

### Cell culture parasite competitions in HFFs, MEFs, RAW264.7, and BMDMs

Competitions were conducted with a red WT strain and green KO or complemented strains. Intracellular parasites were scraped and mechanically lysed through 27 Ga needles. Parasites were passed through a 5 µM filter to remove host cell debris prior to counting. 8 x 10^5^ parasites of each WT and KO strains were combined, and checked for an approximate 1:1 mix by flow cytometry. After confirmation of the starting ratios, parasites were infected onto HFFs in 1 well of a 6 well plate, and spun at 290 x *g* for 5 min. The competition was passaged in culture for at least 1 passage prior to splitting the population to other host cell contexts. After passage 1, competitions were split to infect HFFs, MEFs, RAW264.7, or CD-1 BMDMs as indicated. In each case, host cells were infected at a low MOI of 0.1 to 0.5. HFFs and BMDMs are contact inhibited or exhibit limited growth in culture, and therefore could be plated days in advance of infection. By contrast, MEF and RAW264.7 cells grow continuously, and were therefore seeded onto 6 wells 24 h prior to infection. At each passage, the ratio of green KO to red WT strains was measured by flow cytometry on a MacsQuant VYB.

### Repeated invasion assay

Repeated invasion assays began as competition experiments as described above. Competitions between WT and *Δrasp1* or complemented parasites were maintained for at least one passage prior to the repeated invasion assay to ensure a stable parasite population. At the start of the invasion assay, wells containing intracellular parasites were washed with PBS. PBS was removed, and 1 ml of DMEM supplemented with 10% IFS was added. Intracellular parasites were then scraped and syringe released. 50 µl of lysed parasites was saved to assay parasite ratios by flow cytometry. 200 µl was taken to infect a coverslip of HFFs to assay primary invasion, and the remainder was infected onto a new confluent HFF 6 well. Plates were spun at 290 x *g* for 5 min to allow parasites to make contact with host cells. After 30 min, the primary invasion coverslips were washed 3x with PBS and fixed with 4% formaldehyde. At the same time, the newly infected 6 well was washed 3x with PBS to remove extracellular parasites. 1 ml of DMEM supplemented with 10% IFS was added to the well, and parasites were force lysed to yield extracellular parasites that had just completed an initial invasion. 50 µl of this pool was measured by flow cytometry to serve as the secondary input. The remainder was placed onto a coverslip and allowed to invade a second host monolayer for 30 min. After 30 min, the wells were washed 3x with PBS, and fixed with 4% formaldehyde. Extracellular parasites were detected by staining coverslips with 1:2000 anti-SAG1 followed by 1:1000 cy5-conjugated anti-mouse secondary. Intracellular parasites were counted based on the fluorescent markers expressed in the WT and KO/complement strains. At least twelve random fields were collected per coverslip, and quantified for presence of intracellular and extracellular parasites of each genotype. The ratio of invaded KO or complemented parasites to invaded WT parasites was then compared to the input ratios obtained by flow cytometry.

### Parasite attenuation in mice

Intracellular parasites maintained in DMEM supplemented with 10% IFS were washed with PBS to remove serum. Intracellular parasites were then scraped in PBS, mechanically released, and filtered through a 5 µM filter. Parasites were then counted and diluted to 1 x 10^5^ parasites/ml. 1 x 10^4^ parasites in 100 µl PBS were then injected I.P. into CD-1 female mice of 8–10 weeks of age. Mouse health was monitored daily after injection until the endpoint of the experiment (day 6 post injection). At least 4 mice were included in each group. Anti-*Toxoplasma* serum ELISAs were conducted to verify infection in surviving mice. Mouse survival plots were generated in Graphpad Prism 9.

### CDCF uptake assay

200,000 parasites were infected onto a 6-well microscopy dish of confluent HFF cells. 30 h post infection, media was removed and wells were washed 1x with PBS. Wells were replenished with 2 ml of Fluorobright without serum. 2.5 µM of CDCFDA was added to each well at time zero. Wells were placed in a 37°C tissue culture incubator. Ten minutes after addition of CDCFDA, media was removed and wells were washed 3x with PBS. Wells were replenished with fresh Fluorobright media and immediately imaged for a 10 min time point. After imaging, wells were returned to the 37°C incubator for an additional 20 min. After this incubation, wells were imaged again for a 30 min time point. Vacuoles were quantified for permeability to the metabolized product (CDCF) produced in the host cytosol as well as for the presence of normal or expanded vacuoles.

### Transmission electron microscopy on WT, *Δgra72*, and GRA72-complement infected host cells

225,000 parasites were infected onto 6-well plates of confluent HFF cells 30 h and 8 h prior to the experimental end point. Sample processing was conducted by two methods conducted independently at the Whitehead Institute (method 1) or Johns Hopkins University (method 2).

#### Method 1

At time of fixation, host cells were lifted from wells with trypsin, quenched in 10 ml of DMEM supplemented with 10% IFS, and spun at 200 x *g* for 3 min at 4°C. Media was aspirated, and cell pellets were resuspended in 1 ml of fixative solution. Fixative solution was made immediately before use by mixing solution A (2% OsO_4_ in dH_2_O) and solution B (2% glutaraldehyde in 100 mM phosphate buffer, pH 6.2) on ice. Cells were fixed on ice for 45 min protected from light. After fixation, cells were pelleted at 200 x *g* for 3 min at 4°C and resuspended in 10 ml of ice-cold 50 mM phosphate buffer. Cells were washed a second time by pelleting at 200 x *g* for 3 min at 4°C and resuspending in 1 ml of ice-cold 50 mM phosphate buffer. Pellets were stored in a 50 mM phosphate buffer at 4°C protected from light until further processing.

#### Method 2

HFFs were infected for 30 h and fixed with 2.5% glutaraldehyde (EM grade, Electron Microscopy Sciences, Hatfield, PA) in 0.1 M sodium cacodylate buffer (pH 7.4) for 1 h at room temperature and processed as previously described^129^. Ultrathin sections of infected cells were stained with osmium tetraoxide before examination with Hitachi 7600 EM under 80 kV.

### BH4 and Folate rescue experiments

*Δgch* (mNeonGreen-positive) and WT (TdTomato-positive) parasites were syringe released and filtered through 5 µm filters. Parasites were then counted and 8 x 10^5^ parasites of each strain were mixed together in DMEM or HPLM. The mixed population was analyzed by flow cytometry to determine the precise starting ratio of parasites. 48 h later, parasites were force lysed, and 100–200 µl were passed to new 6 wells containing either DMEM or HPLM with 10% FBS. For nutrient rescue experiments, wells received either 50 µM BH4 or 30 mg/L folate or the equivalent volume of vehicle (DMSO or water, respectively). 48 h post infection, parasites were syringe released and relative ratios of *Δgch* and WT parasites were determined by flow cytometry on a MACSquant VYB.

### Targeted metabolic screening in DMEM vs. HPLM

A gRNA library targeting all annotated metabolic genes in the *Toxoplasma* genome was cloned as previously described^30^. The library was prepared for transfection by AseI digestion and dialysis. Cas9-expressing parasites were then transfected with the digested library. 24 h post transfection, extracellular parasites were aspirated and plates were washed with PBS. Plates were then replenished with media containing 10 µg/ml DNaseI and 3 µg/ml pyrimethamine. At the first lysis, parasites were split into plates containing DMEM or HPLM supplemented with 10 µg/ml gentamicin and 3% IFS. Parasites were maintained for 3 more passages in these media conditions. Samples of parasites were kept at each passage from each media condition to harvest genomic DNA for subsequent gRNA amplification as previously described. Log_2_(fold changes) of gRNA abundance were calculated relative to input libraries to generate phenotype scores of parasites passaged in DMEM and HPLM (**Table S9**).

### Pyrimethamine/DAHP drug synergy lytic assays in *T. gondii*

Eight-point dosage curves were generated by treating parasites in 96 well plates to various concentrations of pyrimethamine with and without 2.5 mM DAHP. At the start of the assay, prior to parasite harvest, 96 well plates containing confluent fibroblasts were set up with 1.33x the desired final drug concentrations in 150 µl of volume. This ensured that the addition of 50ul of parasites would result in the desired 1x drug concentration of both pyrimethamine and DAHP. Wells designated as containing less than the maximum dosage of either drug (10 µg/ml pyrimethamine and 2.5 mM DAHP) received the equivalent volume of vehicle (ethanol for pyrimethamine, PBS for DAHP). Parasites were then mechanically-lysed, filtered, and resuspended to a concentration of 5 x 10^5^ parasites/ml. 50 µl of parasites were then added to each well, resulting in an infection of 2.5 x 10^4^ parasites per well. Plates were then spun at 290 *g* for 5 min. Plates were then incubated for 3 days to allow for parasite plaquing. Plates were then washed with PBS followed by staining with crystal violet as described for plaque assays above. Staining was then quantified by measuring absorbance at 570 nm. Percent lysis for each drug condition was normalized to wells containing parasites with no drugs (minimum staining) and as well wells containing no parasites (maximum staining). Drug curves were then generated and IC_50_ values were calculated from biological triplicate experiments in GraphPad Prism 9.

### Dose-response assay in *Plasmodium falciparum*

A SYBR Green I–based assay was used to measure *in vitro P. falciparum* drug susceptibility^168, 169^. Ring-stage NF54 parasites were seeded at 1% hematocrit and 1% starting parasitemia in 384-well black clear-bottom plates containing test compounds (stocks of 10 mM pyrimethamine, 0.1 mM atovaquone, and 10 mM DHA in DMSO) plated in triplicate in 12-point serial dilutions for 72h. The parasites were plated in complete RPMI 1640 media supplemented with 25 mM HEPES, 0.21% sodium bicarbonate, 50 mg/l hypoxanthine, and 0.5% Albumax II (Invitrogen) with or without 1 mM DAHP. Lysis buffer (0.16% w/v saponin, 1.6% Triton X-100, 5 mM EDTA, and 20 mM Tris-HCl, pH 7.4) with a 1:1000 dilution of SYBR Green I fluorescent dye (Invitrogen) was added for at least 12 h, and fluorescence readings were taken (excitation at 494 nm, emission at 530 nm) using a SpectraMax M5 (Molecular Devices, Sunnyvale, CA) plate reader. IC_50_ values were calculated using a nonlinear regression curve fit in Graphpad Prism 9 in biological triplicate.

### Statistical analysis and Graphing

Graphs were generated in Graphpad Prism 9 or in R using ggplot or other packages indicated in the relevant methods section.

## ACKNOWLEDGEMENTS

We thank Christopher A. Hunter, Anita A. Koshy, Jeroen P. Saeij, and all members of the Lourido Lab for helpful discussions and support. Emily Shortt additionally provided key technical support. We thank Jeroen P. Saeij for the *Δgra17* and GRA17-complemented strains, Louis M. Weiss for the anti-MAG1 antibody, David W. Threadgill for the CAST/EiJ and C57Bl/6J MEF cell lines, Bryan D. Bryson for the RAW264.7 cell line, Peter J. Bradley for the anti-GRA7 and anti-ROP13 antibodies, L. David Sibley for hybridomas for anti-TY and anti-SAG1 antibodies. Wandy L. Beatty at the Molecular Microbiology Imaging facility in Washington University performed the sample processing and transmission electron microscopy for initial analysis of the *Δgra72*. We thank the technical staff of the Electron Microscopy Core Facility at the Johns Hopkins University School of Medicine for assistance with secondary analysis *Δgra72* by transmission electron microscopy. This work relied extensively on VEuPathDB.org and we thank all contributors to this resource. This work was supported by the Burroughs Wellcome Fund PATH award and grants from the National Institutes of Health (R01 AI158501 and R01 AI144369) to S.L., and a National Science Foundation GRFP fellowship to C.J.G..

## AUTHOR CONTRIBUTIONS

C.J.G., S.L., and J.D.D. conceptualized this project. C.J.G., K.J.W., F.M.H., B.S.W, M.F., E.A.B., T.C.T.L., A.L.H., A.G.S., and I.C., performed all experiments. C.J.G., K.J.W., and R.A.T performed all data analysis. C.J.G. and S.L. wrote the manuscript with input from all authors.

## DATA AVAILABILITY

All screen data is available in supplementary tables. UMI barcode counts are available upon reasonable request, and will be deposited in a publicly accessible database prior to publication. Materials including strains and plasmids are available upon reasonable request.

## CODE AVAILABILITY

All code is available upon reasonable request.

---

**Figure S1.**
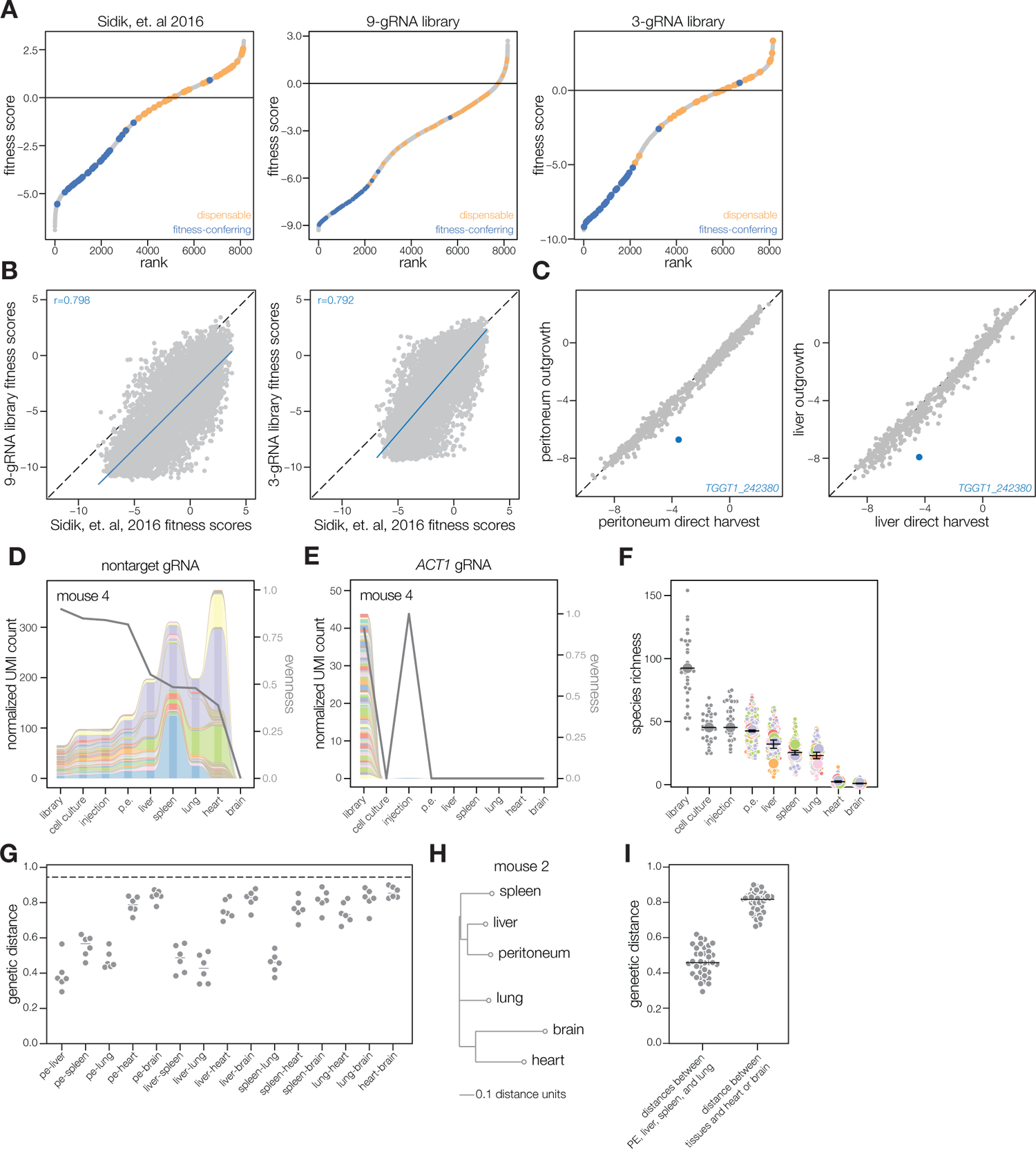
**(A)** Rank-ordered plots of gene fitness scores from a genome-wide CRISPR screen published in Sidik, et. al 2016, or screens using new nine-gRNA and three-gRNA libraries. Control dispensable and fitness-conferring genes are highlighted in orange and blue, respectively. **(B)** Scatter plots comparing gene fitness scores from nine-gRNA or three-gRNA library screens against fitness scores from Sidik, et. al^10^. Linear regressions and Pearson correlation scores are colored in blue. **(C)** Scatter plots comparing gene fitness scores from screen samples harvested directly from peritoneum or liver against those derived from parasite outgrowth from infected tissues. *TGGT1_242380*, a gene identified as fitness-conferring in parasite infection of bone-marrow derived macrophages is highlighted in blue^13^. **(D)** Alluvial plots depicting relative barcode abundance of a nontargeting gRNA across tissues in a representative mouse. Line graph indicates evenness of barcodes in each sample. **(E)** Alluvial plots depicting relative barcode abundance of a *ACT1* targeting gRNA across tissues in a representative mouse. Line graph indicates evenness of barcodes in each sample. **(F)** Superplot of species richness for nontargeting gRNAs across tissues from six mice. Colors indicate different mice with small points for each gRNA and large points for the mean of all gRNAs in each mouse. Mean ± SEM for all mice is overplotted. **(G)** Genetic distance of barcode populations between indicated tissues across six mice. **(H)** Dendrograms of tissues based on genetic distance of barcode relatedness from nontargeting gRNAs. **(I)** Average genetic distances between peritoneum, liver, spleen, and lung compared to average distances between tissues to heart and brain.

**Figure S2.**
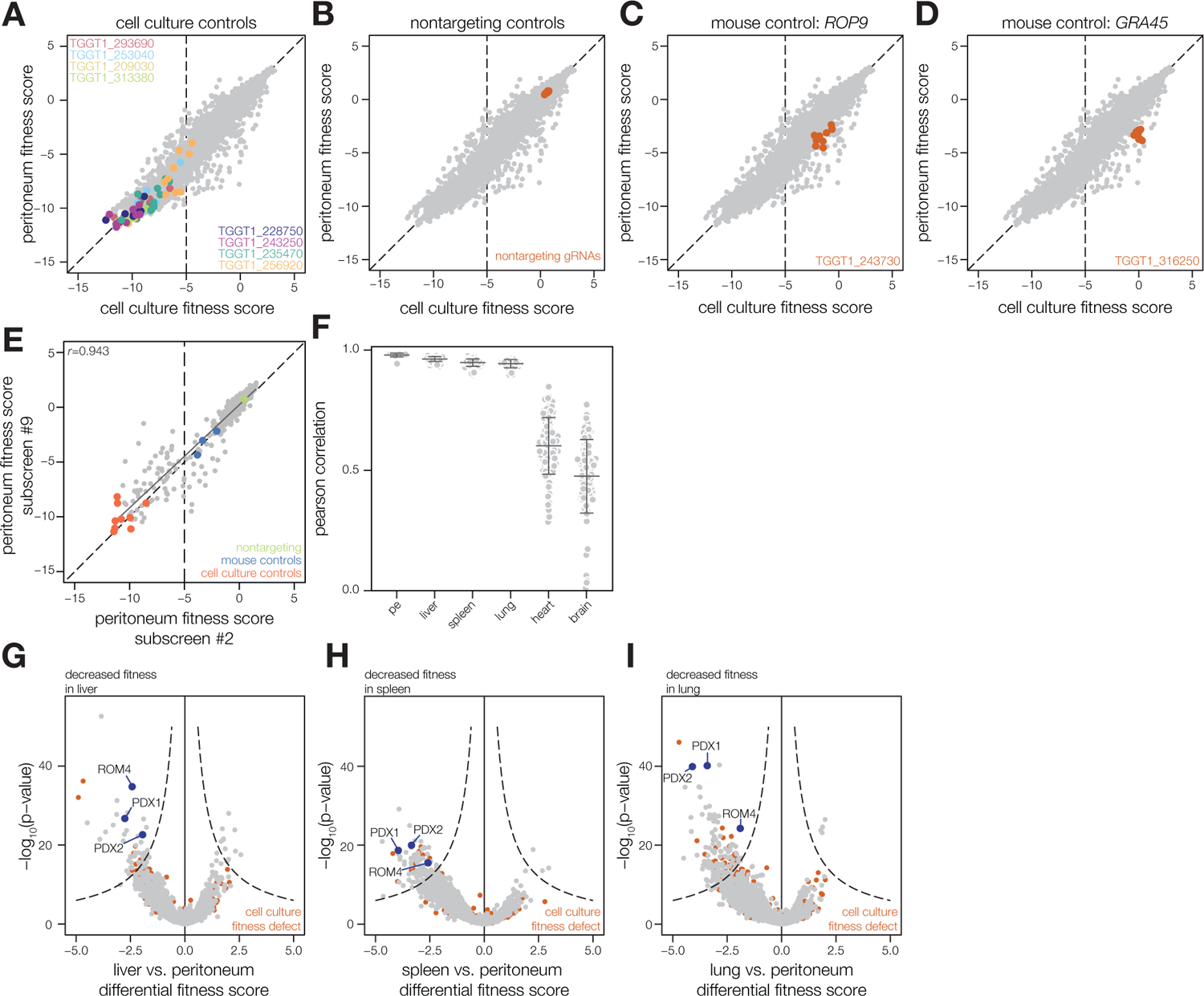
**(A–D)** Scatter plot comparing gene fitness scores from nine subscreens from peritoneum and cell culture (passage 8), highlighting cell culture essential controls **(A)**, nontargeting controls **(B)**, or two control genes required for mouse infection, *ROP9* **(C)**, and *GRA45* **(D)**. **(E)** Scatter plot comparing gene fitness scores from sublibrary three screened in subscreen two and subscreen nine (Pearson correlation *r* = 0.943). **(F)** Pearson correlations of gene fitness scores between mice within a subscreen for each tissue. Mean ± SD indicated. **(G-I)** Volcano plots of differential fitness of liver, spleen, or lung relative to peritoneum against - log_10_(p-value). Dotted line indicates a threshold of [-log_10_(pval) · differential fitness score] = 30. Genes with cell culture fitness scores <-5 are colored in red.

**Figure S3.**
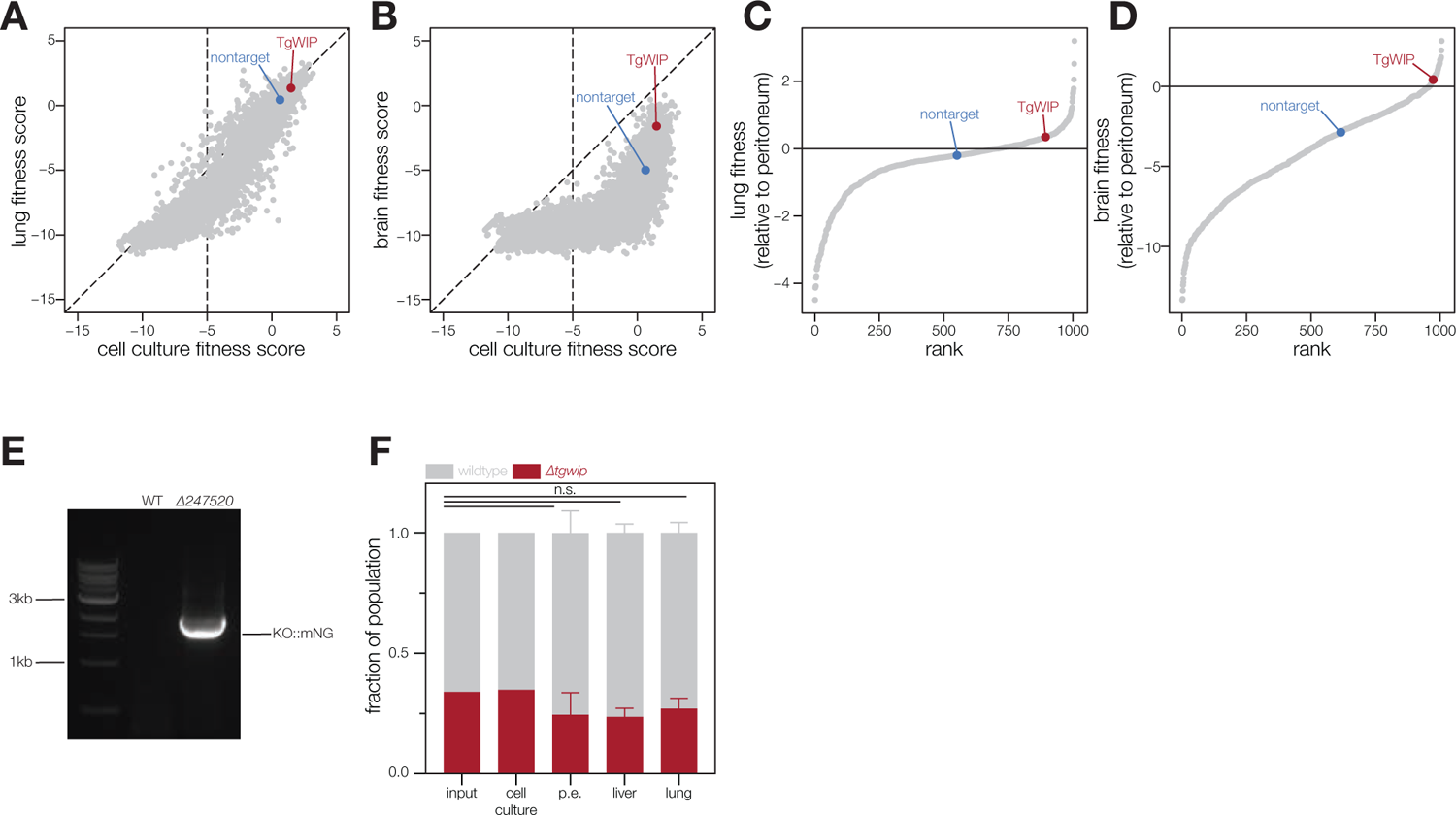
**(A–B)** Scatter plot comparing gene fitness scores from lung or brain to cell culture (passage 8). *TgWIP* and nontargeting control fitness scores are highlighted in red and blue, respectively. **(C-D)** Rank-ordered plot of gene fitness scores generated by normalization to peritoneum abundance for lung and brain. *TgWIP* and nontargeting control fitness scores are highlighted in red and blue, respectively. **(E)** DNA gel depicting *TgWip* knockout generated by replacing the locus with a mNeonGreen fluorophore. **(F)** Fraction of *Δtgwip* and WT parasites from the indicated sample during a co-injection competition experiment. Mean ± SD indicated (*n* = 3 mice for each tissue, unpaired two-tailed *t*-test).

**Figure S4.**
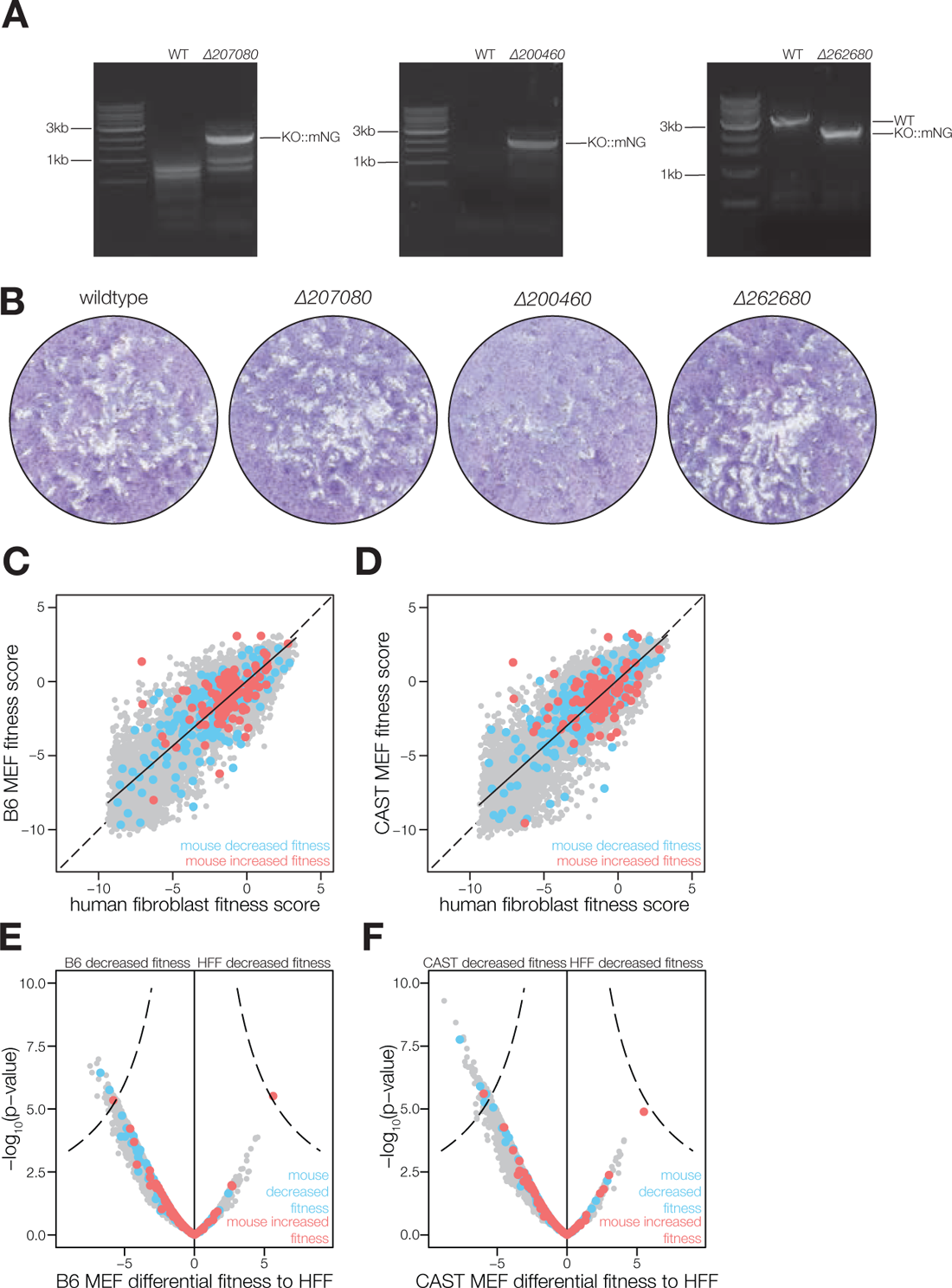
**(A)** DNA gels depicting knockout generation by replacing the indicated loci with a mNeonGreen fluorophore. **(B)** Plaque assays from 7 days of growth of 500 parasites of the indicated genotype. **(C-D)** Scatter plots comparing gene fitness scores from parasites cultured in B6 or CAST MEF cell lines compared to parasites cultured in HFFs. Candidates with decreased and increased fitness in the mouse peritoneum are highlighted in blue and red, respectively. **(E-F)** Volcano plot of differential fitness between gene scores from B6 or CAST MEFs relative to HFF against -log_10_(p-value). Dotted line indicates a threshold of [-log_10_(pval) · differential fitness score] = 30. Candidates with decreased and increased fitness in the mouse screen are highlighted in blue and red, respectively.

**Figure S5.**
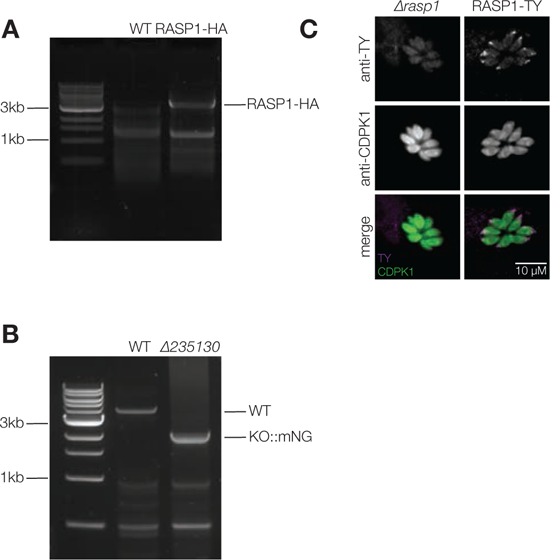
**(A)** DNA gel depicting integration of an HA tag at the endogenous *RASP1* (*TGGT1_235130*) locus. **(B)** DNA gel depicting RASP1 knockout generated by replacing the locus with a mNeonGreen fluorophore. **(C)** IFA of RASP1-TY complement strain with CDPK1 counterstain.

**Figure S6.**
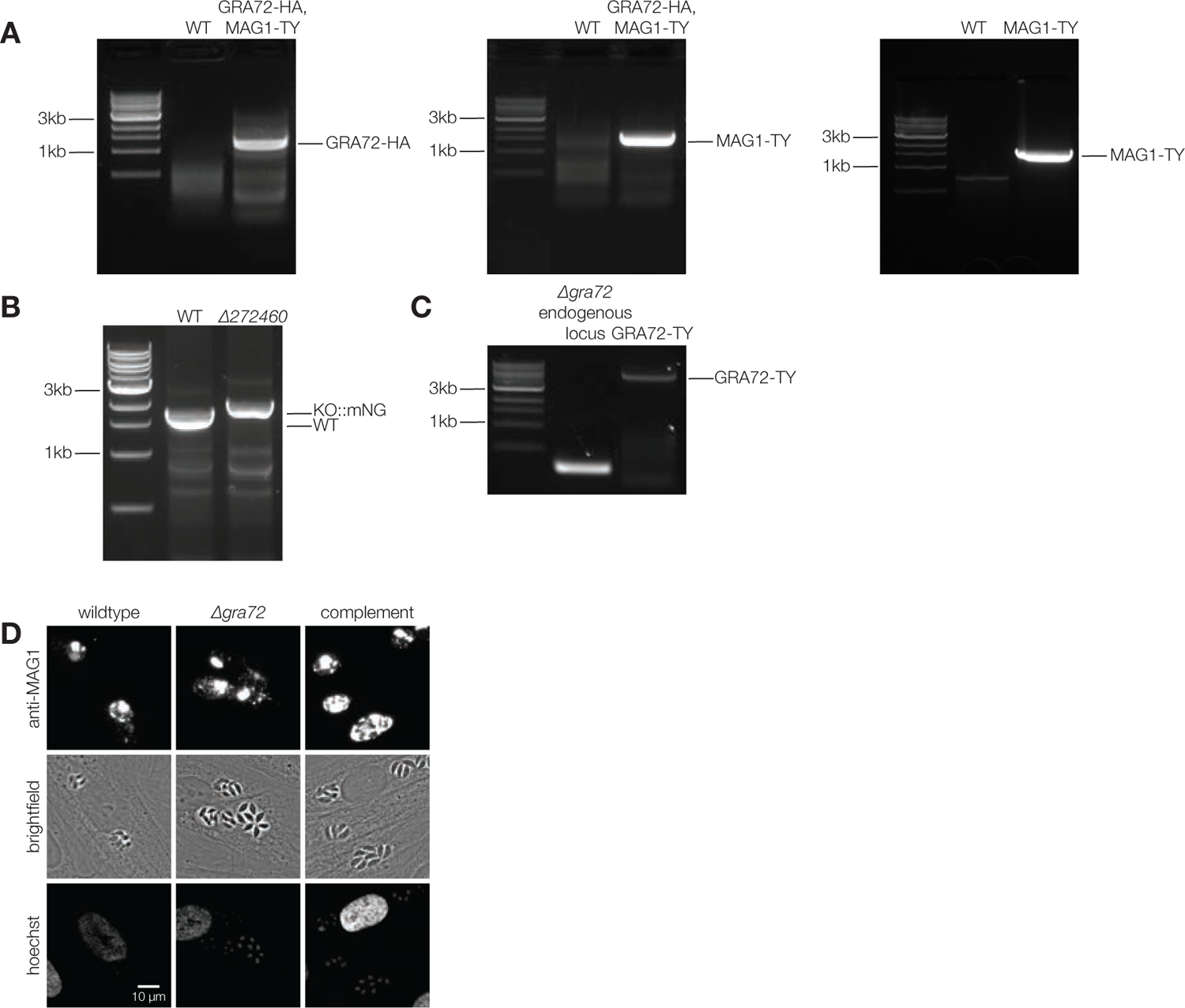
**(A)** DNA gel depicting integration of an HA tag at the endogenous *GRA72* (*TGGT1_272460*) locus and a TY tag at the endogenous *MAG1* (*TGGT1_270240*) locus in MAG1-TY/GRA72-HA and MAG1-TY strains. **(B)** DNA gel depicting GRA72 knockout generated by replacing the locus with a mNeonGreen fluorophore. **(C)** DNA gel depicting integration of GRA72-TY construct into an intergenic locus in the *Δgra72* parental strain. **(D)** IFA of MAG1 from WT, *Δgra72*, or complemented parasites at 24 h post infection.

**Figure S7.**
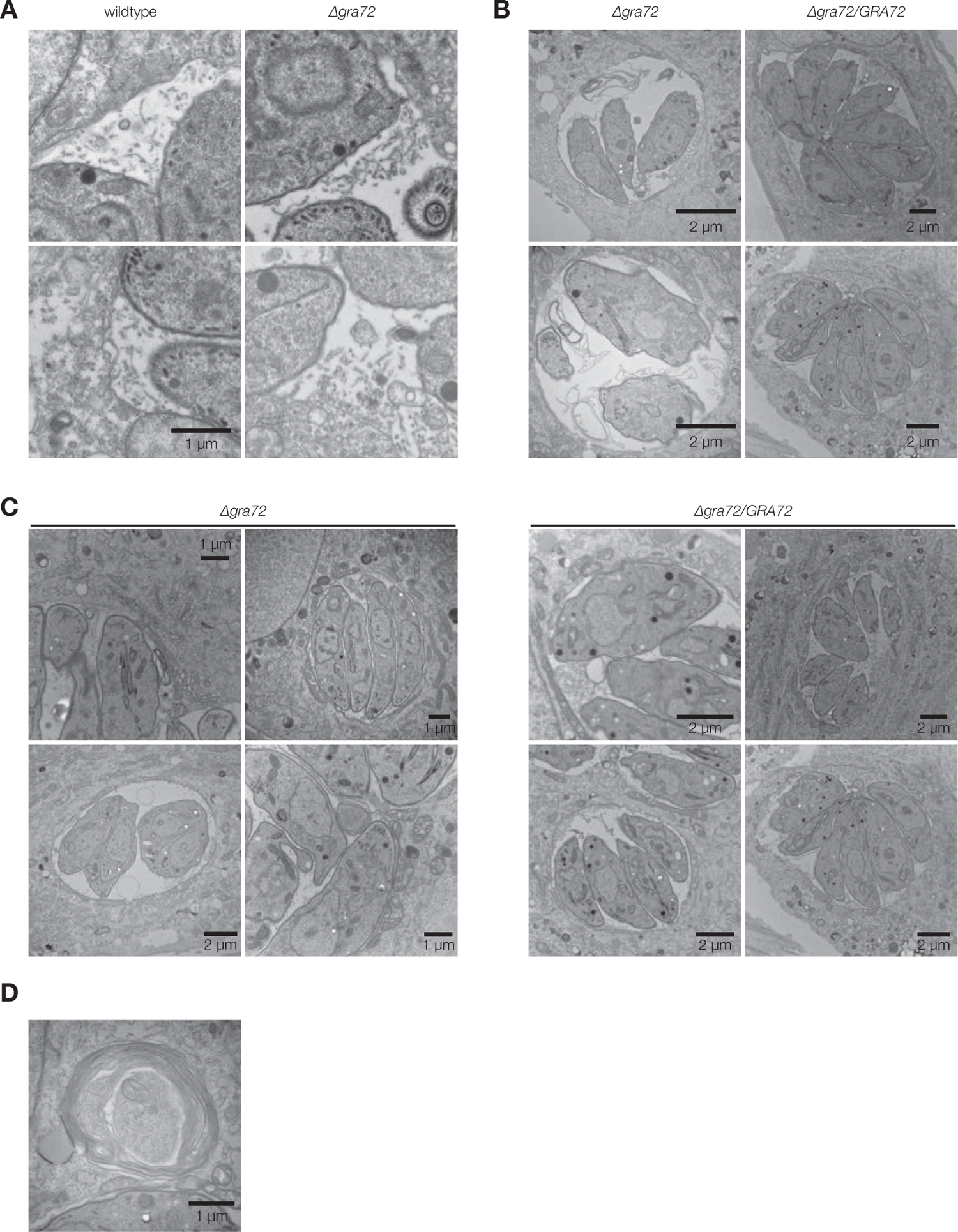
**(A)** Electron Micrograph depicting intravacuolar network in WT and *Δgra72* parasite vacuoles. **(B)** Electron Micrograph depicting vacuole membrane irregularities in *Δgra72* parasite vacuoles. **(C)** Electron Micrograph depicting host organelle recruitment (mitochondria, ER, and golgi) by *Δgra72* and *Δgra72/GRA72-TY* vacuoles. **(D)** Electron Micrograph depicting host material within a developing multi-lamellar body.

**Figure S8.**
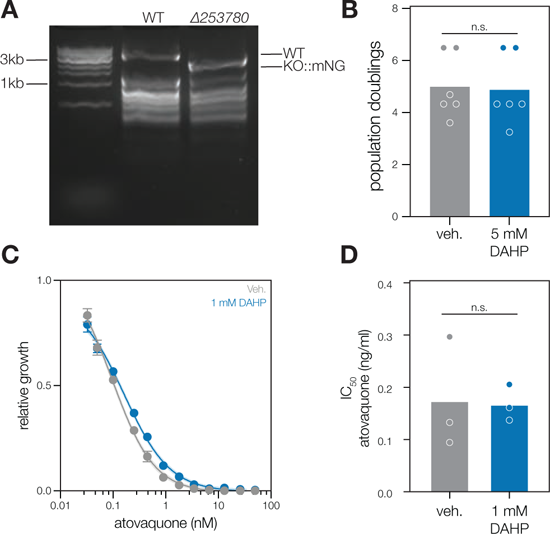
**(A)** DNA gel depicting GCH knockout generated by replacing the locus with a mNeonGreen fluorophore. **(B)** Population doubling of C57BL/6J immortalized MEFs in the presence of PBS vehicle or 5 mM DAHP (*n* = 6, unpaired two-tailed *t*-test). **(C)** 12-point dose response curve of *P. falciparum* parasites grown in varying concentrations of atovaquone with and without 1 mM DAHP co-treatment. **(D)** IC_50_ value of atovaquone for *P. falciparum* parasites with and without DAHP co-treatment (*n* = 3, unpaired two-tailed *t*-test).

## REFERENCES

1. Pasqual, G. et al. Monitoring T cell-dendritic cell interactions in vivo by intercellular enzymatic labelling. Nature 553, 496–500 (2018).

2. Chan, M. M. et al. Molecular recording of mammalian embryogenesis. Nature 570, 77–82 (2019).

3. Bowling, S. et al. An Engineered CRISPR-Cas9 Mouse Line for Simultaneous Readout of Lineage Histories and Gene Expression Profiles in Single Cells. Cell 181, 1410–1422.e27 (2020).

4. Rodriques, S. G. et al. Slide-seq: A scalable technology for measuring genome-wide expression at high spatial resolution. Science 363, 1463–1467 (2019).

5. Montoya, J. G. & Liesenfeld, O. Toxoplasmosis. Lancet 363, 1965–1976 (2004).

6. Sibley, L. D., Khan, A., Ajioka, J. W. & Rosenthal, B. M. Genetic diversity of Toxoplasma gondii in animals and humans. Philos. Trans. R. Soc. Lond. B Biol. Sci. 364, 2749–2761 (2009).

7. Lawley, T. D. et al. Genome-wide screen for Salmonella genes required for long-term systemic infection of the mouse. PLoS Pathog. 2, e11 (2006).

8. Keys, H. R. & Knouse, K. A. Genome-scale CRISPR screening in a single mouse liver. Cell Genomics 0, (2022).

9. Chen, S. et al. Genome-wide CRISPR screen in a mouse model of tumor growth and metastasis. Cell 160, 1246–1260 (2015).

10. Sidik, S. M. et al. A Genome-wide CRISPR Screen in Toxoplasma Identifies Essential Apicomplexan Genes. Cell 166, 1423–1435.e12 (2016).

11. Harding, C. R. et al. Genetic screens reveal a central role for heme metabolism in artemisinin susceptibility. Nat. Commun. 11, 4813 (2020).

12. Chen, Y. et al. Genome-Wide CRISPR/Cas9 Screen Identifies New Genes Critical for Defense Against Oxidant Stress in Toxoplasma gondii. Front. Microbiol. 12, 670705 (2021).

13. Wang, Y. et al. Genome-wide screens identify Toxoplasma gondii determinants of parasite fitness in IFNγ-activated murine macrophages. Nat. Commun. 11, 5258 (2020).

14. Sangaré, L. O. et al. In Vivo CRISPR Screen Identifies TgWIP as a Toxoplasma Modulator of Dendritic Cell Migration. Cell Host Microbe 26, 478–492.e8 (2019).

15. Young, J. et al. A CRISPR platform for targeted in vivo screens identifies Toxoplasma gondii virulence factors in mice. Nat. Commun. 10, 3963 (2019).

16. Butterworth, S., et al. Toxoplasma gondii ROP1 subverts murine and human innate immune restriction. *bioRxiv* 2022.03.21.485090 (2022) doi:10.1101/2022.03.21.485090.

17. Bushell, E. et al. Functional Profiling of a Plasmodium Genome Reveals an Abundance of Essential Genes. Cell 170, 260–272.e8 (2017).

18. Michlits, G. et al. CRISPR-UMI: single-cell lineage tracing of pooled CRISPR-Cas9 screens. Nat. Methods 14, 1191–1197 (2017).

19. Wong, A. S. L. et al. Multiplexed barcoded CRISPR-Cas9 screening enabled by CombiGEM. Proc. Natl. Acad. Sci. U. S. A. 113, 2544–2549 (2016).

20. Zhu, S. et al. Guide RNAs with embedded barcodes boost CRISPR-pooled screens. Genome Biol. 20, 20 (2019).

21. Schmierer, B. et al. CRISPR/Cas9 screening using unique molecular identifiers. Mol. Syst. Biol. 13, 945 (2017).

22. Doench, J. G. et al. Optimized sgRNA design to maximize activity and minimize off-target effects of CRISPR-Cas9. Nat. Biotechnol. 34, 184–191 (2016).

23. Djurković-Djaković, O. et al. Kinetics of parasite burdens in blood and tissues during murine toxoplasmosis. Exp. Parasitol. 131, 372–376 (2012).

24. Andenmatten, N. et al. Conditional genome engineering in Toxoplasma gondii uncovers alternative invasion mechanisms. Nat. Methods 10, 125–127 (2013).

25. Drewry, L. L. & Sibley, L. D. Toxoplasma Actin Is Required for Efficient Host Cell Invasion. MBio 6, e00557 (2015).

26. Behnke, M. S. et al. The polymorphic pseudokinase ROP5 controls virulence in Toxoplasma gondii by regulating the active kinase ROP18. PLoS Pathog. 8, e1002992 (2012).

27. Etheridge, R. D. et al. The Toxoplasma pseudokinase ROP5 forms complexes with ROP18 and ROP17 kinases that synergize to control acute virulence in mice. Cell Host Microbe 15, 537–550 (2014).

28. Behnke, M. S. et al. Virulence differences in Toxoplasma mediated by amplification of a family of polymorphic pseudokinases. Proc. Natl. Acad. Sci. U. S. A. 108, 9631–9636 (2011).

29. Konrad, C., Wek, R. C. & Sullivan, W. J., Jr. A GCN2-like eukaryotic initiation factor 2 kinase increases the viability of extracellular Toxoplasma gondii parasites. Eukaryot. Cell 10, 1403–1412 (2011).

30. Krishnan, A. et al. Functional and Computational Genomics Reveal Unprecedented Flexibility in Stage-Specific Toxoplasma Metabolism. Cell Host Microbe 27, 290–306.e11 (2020).

31. Krishnan, A., Kloehn, J., Lunghi, M. & Soldati-Favre, D. Vitamin and cofactor acquisition in apicomplexans: Synthesis versus salvage. J. Biol. Chem. 295, 701–714 (2020).

32. Chattopadhyay, R. et al. PfSPATR, a Plasmodium falciparum protein containing an altered thrombospondin type I repeat domain is expressed at several stages of the parasite life cycle and is the target of inhibitory antibodies. J. Biol. Chem. 278, 25977–25981 (2003).

33. Huynh, M.-H., Boulanger, M. J. & Carruthers, V. B. A conserved apicomplexan microneme protein contributes to Toxoplasma gondii invasion and virulence. Infect. Immun. 82, 4358– 4368 (2014).

34. Kafsack, B. F. C. et al. Rapid membrane disruption by a perforin-like protein facilitates parasite exit from host cells. Science 323, 530–533 (2009).

35. Dongchao, Z., Ning, J. & Qijun, C. Loss of rhoptry protein 9 impeded Toxoplasma gondii infectivity. Acta Trop. 207, 105464 (2020).

36. Peixoto, L. et al. Integrative genomic approaches highlight a family of parasite-specific kinases that regulate host responses. Cell Host Microbe 8, 208–218 (2010).

37. Michelin, A. et al. GRA12, a Toxoplasma dense granule protein associated with the intravacuolar membranous nanotubular network. Int. J. Parasitol. 39, 299–306 (2009).

38. Fox, B. A. et al. Toxoplasma gondii Parasitophorous Vacuole Membrane-Associated Dense Granule Proteins Orchestrate Chronic Infection and GRA12 Underpins Resistance to Host Gamma Interferon. MBio 10, (2019).

39. Coffey, M. J. et al. Aspartyl Protease 5 Matures Dense Granule Proteins That Reside at the Host-Parasite Interface in Toxoplasma gondii. MBio 9, (2018).

40. Shastri, A. J., Marino, N. D., Franco, M., Lodoen, M. B. & Boothroyd, J. C. GRA25 is a novel virulence factor of Toxoplasma gondii and influences the host immune response. Infect. Immun. 82, 2595–2605 (2014).

41. Franco Magdalena et al. A Novel Secreted Protein, MYR1, Is Central to Toxoplasma’s Manipulation of Host Cells. MBio 7, e02231–15 (2016).

42. Marino, N. D. et al. Identification of a novel protein complex essential for effector translocation across the parasitophorous vacuole membrane of Toxoplasma gondii. PLoS Pathog. 14, e1006828 (2018).

43. Cygan, A. M. et al. Coimmunoprecipitation with MYR1 Identifies Three Additional Proteins within the Toxoplasma gondii Parasitophorous Vacuole Required for Translocation of Dense Granule Effectors into Host Cells. mSphere 5, (2020).

44. Harker, K. S., Ueno, N. & Lodoen, M. B. Toxoplasma gondii dissemination: a parasite’s journey through the infected host. Parasite Immunol. 37, 141–149 (2015).

45. Buguliskis, J. S., Brossier, F., Shuman, J. & Sibley, L. D. Rhomboid 4 (ROM4) affects the processing of surface adhesins and facilitates host cell invasion by Toxoplasma gondii. PLoS Pathog. 6, e1000858 (2010).

46. Kenthirapalan, S., Waters, A. P., Matuschewski, K. & Kooij, T. W. A. Copper-transporting ATPase is important for malaria parasite fertility. Molecular Microbiology vol. 91 315–325 Preprint at https://doi.org/10.1111/mmi.12461 (2014).

47. Xue, J. et al. Thioredoxin reductase from Toxoplasma gondii: an essential virulence effector with antioxidant function. FASEB J. 31, 4447–4457 (2017).

48. Rahman, K. et al. The E3 Ubiquitin Ligase Adaptor Protein Skp1 Is Glycosylated by an Evolutionarily Conserved Pathway That Regulates Protist Growth and Development. Journal of Biological Chemistry vol. 291 4268–4280 Preprint at https://doi.org/10.1074/jbc.m115.703751 (2016).

49. West, C. M. & Blader, I. J. Oxygen sensing by protozoans: how they catch their breath. Curr. Opin. Microbiol. 26, 41–47 (2015).

50. Primo, V. A., Jr, et al. The Extracellular Milieu of Toxoplasma’s Lytic Cycle Drives Lab Adaptation, Primarily by Transcriptional Reprogramming. mSystems 6, e0119621 (2021).

51. Mas-Bargues, C., et al. Relevance of Oxygen Concentration in Stem Cell Culture for Regenerative Medicine. Int. J. Mol. Sci. 20, (2019).

52. Barylyuk, K. et al. A Comprehensive Subcellular Atlas of the Toxoplasma Proteome via hyperLOPIT Provides Spatial Context for Protein Functions. Cell Host Microbe 28, 752– 766.e9 (2020).

53. Vonlaufen, N., Naguleswaran, A., Coppens, I. & Sullivan, W. J., Jr. MYST family lysine acetyltransferase facilitates ataxia telangiectasia mutated (ATM) kinase-mediated DNA damage response in Toxoplasma gondii. J. Biol. Chem. 285, 11154–11161 (2010).

54. Moreira-Souza, A. C. A. et al. The P2X7 Receptor Mediates Toxoplasma gondii Control in Macrophages through Canonical NLRP3 Inflammasome Activation and Reactive Oxygen Species Production. Front. Immunol. 8, 1257 (2017).

55. Zhuang, H. et al. DNA double-strand breaks in the Toxoplasma gondii-infected cells by the action of reactive oxygen species. Parasit. Vectors 13, 490 (2020).

56. Zhang, H. et al. Toxoplasma gondii UBL-UBA shuttle proteins contribute to the degradation of ubiquitinylated proteins and are important for synchronous cell division and virulence. FASEB J. 34, 13711–13725 (2020).

57. Boothroyd, J. C. & Dubremetz, J.-F. Kiss and spit: the dual roles of Toxoplasma rhoptries. Nat. Rev. Microbiol. 6, 79–88 (2008).

58. Suarez, C. et al. A lipid-binding protein mediates rhoptry discharge and invasion in Plasmodium falciparum and Toxoplasma gondii parasites. Nat. Commun. 10, 4041 (2019).

59. Liffner, B. et al. PfCERLI1 is a conserved rhoptry associated protein essential for Plasmodium falciparum merozoite invasion of erythrocytes. Nat. Commun. 11, 1411 (2020).

60. Liffner, B. et al. Cell biological analysis reveals an essential role for Pfcerli2 in erythrocyte invasion by malaria parasites. Commun Biol 5, 121 (2022).

61. Markus, B. M., Waldman, B. S., Lorenzi, H. A. & Lourido, S. High-Resolution Mapping of Transcription Initiation in the Asexual Stages of Toxoplasma gondii. Front. Cell. Infect. Microbiol. 10, 617998 (2020).

62. Park, J. & Hunter, C. A. The role of macrophages in protective and pathological responses to Toxoplasma gondii. Parasite Immunol. 42, e12712 (2020).

63. Delgado Betancourt, E., et al. From Entry to Early Dissemination-Toxoplasma gondii’s Initial Encounter With Its Host. Front. Cell. Infect. Microbiol. 9, 46 (2019).

64. Courret, N. et al. CD11c- and CD11b-expressing mouse leukocytes transport single Toxoplasma gondii tachyzoites to the brain. Blood 107, 309–316 (2006).

65. Mageswaran, S. K. et al. In situ ultrastructures of two evolutionarily distant apicomplexan rhoptry secretion systems. Nat. Commun. 12, 4983 (2021).

66. Chen, X.-M. et al. Apical organelle discharge by Cryptosporidium parvum is temperature, cytoskeleton, and intracellular calcium dependent and required for host cell invasion. Infect. Immun. 72, 6806–6816 (2004).

67. Koshy, A. A. et al. Toxoplasma co-opts host cells it does not invade. PLoS Pathog. 8, e1002825 (2012).

68. Tomita, T. et al. Toxoplasma gondii Matrix Antigen 1 Is a Secreted Immunomodulatory Effector. MBio 12, (2021).

69. Gold, D. A. et al. The Toxoplasma Dense Granule Proteins GRA17 and GRA23 Mediate the Movement of Small Molecules between the Host and the Parasitophorous Vacuole. Cell Host Microbe 17, 642–652 (2015).

70. Wang, J.-L. et al. Immunization with Toxoplasma gondii GRA17 Deletion Mutant Induces Partial Protection and Survival in Challenged Mice. Front. Immunol. 8, 730 (2017).

71. Paredes-Santos, T., Wang, Y., Waldman, B., Lourido, S. & Saeij, J. P. The GRA17 Parasitophorous Vacuole Membrane Permeability Pore Contributes to Bradyzoite Viability. Front. Cell. Infect. Microbiol. 9, 321 (2019).

72. Alaganan, A., Fentress, S. J., Tang, K., Wang, Q. & Sibley, L. D. *Toxoplasma* GRA7 effector increases turnover of immunity-related GTPases and contributes to acute virulence in the mouse. Proc. Natl. Acad. Sci. U. S. A. 111, 1126–1131 (2014).

73. Dunn, J. D., Ravindran, S., Kim, S.-K. & Boothroyd, J. C. The Toxoplasma gondii dense granule protein GRA7 is phosphorylated upon invasion and forms an unexpected association with the rhoptry proteins ROP2 and ROP4. Infect. Immun. 76, 5853–5861 (2008).

74. Panas, M. W. & Boothroyd, J. C. Toxoplasma Uses GRA16 To Upregulate Host c-Myc. mSphere 5, (2020).

75. Sibley, L. D., Niesman, I. R., Parmley, S. F. & Cesbron-Delauw, M. F. Regulated secretion of multi-lamellar vesicles leads to formation of a tubulo-vesicular network in host-cell vacuoles occupied by Toxoplasma gondii. J. Cell Sci. 108 **(Pt** **4**), 1669–1677 (1995).

76. Romano, J. D. et al. The parasite Toxoplasma sequesters diverse Rab host vesicles within an intravacuolar network. J. Cell Biol. 216, 4235–4254 (2017).

77. Sinai, A. P., Webster, P. & Joiner, K. A. Association of host cell endoplasmic reticulum and mitochondria with the Toxoplasma gondii parasitophorous vacuole membrane: a high affinity interaction. J. Cell Sci. 110 **(Pt** **17**), 2117–2128 (1997).

78. Melo, E. J. & de Souza, W. Relationship between the host cell endoplasmic reticulum and the parasitophorous vacuole containing Toxoplasma gondii. Cell Struct. Funct. 22, 317– 323 (1997).

79. Pernas, L. et al. Toxoplasma effector MAF1 mediates recruitment of host mitochondria and impacts the host response. PLoS Biol. 12, e1001845 (2014).

80. Deffieu, M. S., Alayi, T. D., Slomianny, C. & Tomavo, S. The Toxoplasma gondii dense granule protein TgGRA3 interacts with host Golgi and dysregulates anterograde transport. Biol. Open 8, (2019).

81. Romano, J. D., Sonda, S., Bergbower, E., Smith, M. E. & Coppens, I. Toxoplasma gondii salvages sphingolipids from the host Golgi through the rerouting of selected Rab vesicles to the parasitophorous vacuole. Mol. Biol. Cell 24, 1974–1995 (2013).

82. Coppens, I. & Romano, J. D. Hostile intruder: Toxoplasma holds host organelles captive. PLoS Pathog. 14, e1006893 (2018).

83. Hariri, M. et al. Biogenesis of multilamellar bodies via autophagy. Mol. Biol. Cell 11, 255– 268 (2000).

84. Ogata, M. et al. Autophagy is activated for cell survival after endoplasmic reticulum stress. Mol. Cell. Biol. 26, 9220–9231 (2006).

85. Coppens, I., Sinai, A. P. & Joiner, K. A. Toxoplasma gondii exploits host low-density lipoprotein receptor-mediated endocytosis for cholesterol acquisition. J. Cell Biol. 149, 167–180 (2000).

86. Romano, J. D., et al. Toxoplasma scavenges mammalian host organelles through usurpation of host ESCRT-III and Vps4. *bioRxiv* 2022.04.11.487923 (2022) doi:10.1101/2022.04.11.487923.

87. Pernas, L., Bean, C., Boothroyd, J. C. & Scorrano, L. Mitochondria Restrict Growth of the Intracellular Parasite Toxoplasma gondii by Limiting Its Uptake of Fatty Acids. Cell Metab. 27, 886–897.e4 (2018).

88. Tymoshenko, S. et al. Metabolic Needs and Capabilities of Toxoplasma gondii through Combined Computational and Experimental Analysis. PLoS Comput. Biol. 11, e1004261 (2015).

89. Blume, M. & Seeber, F. Metabolic interactions between Toxoplasma gondii and its host. F1000Res. 7, (2018).

90. Mazumdar, J. & Striepen, B. Make it or take it: fatty acid metabolism of apicomplexan parasites. Eukaryot. Cell 6, 1727–1735 (2007).

91. Hartmann, A., Hellmund, M., Lucius, R., Voelker, D. R. & Gupta, N. Phosphatidylethanolamine synthesis in the parasite mitochondrion is required for efficient growth but dispensable for survival of Toxoplasma gondii. J. Biol. Chem. 289, 6809–6824 (2014).

92. Müller, S. & Kappes, B. Vitamin and cofactor biosynthesis pathways in Plasmodium and other apicomplexan parasites. Trends Parasitol. 23, 112–121 (2007).

93. Yang, J., He, Z., Chen, C., Zhao, J. & Fang, R. Starch Branching Enzyme 1 Is Important for Amylopectin Synthesis and Cyst Reactivation in Toxoplasma gondii. Microbiol Spectr 10, e0189121 (2022).

94. Cantor, J. R. et al. Physiologic Medium Rewires Cellular Metabolism and Reveals Uric Acid as an Endogenous Inhibitor of UMP Synthase. Cell vol. 169 258–272.e17 Preprint at https://doi.org/10.1016/j.cell.2017.03.023 (2017).

95. Basset, G. et al. Folate synthesis in plants: the first step of the pterin branch is mediated by a unique bimodular GTP cyclohydrolase I. Proc. Natl. Acad. Sci. U. S. A. 99, 12489– 12494 (2002).

96. Kümpornsin, K. et al. Biochemical and functional characterization of Plasmodium falciparum GTP cyclohydrolase I. Malar. J. 13, 150 (2014).

97. Dittrich, S. et al. An atypical orthologue of 6-pyruvoyltetrahydropterin synthase can provide the missing link in the folate biosynthesis pathway of malaria parasites. Mol. Microbiol. 67, 609–618 (2008).

98. Crabtree, M. J. & Channon, K. M. Synthesis and recycling of tetrahydrobiopterin in endothelial function and vascular disease. Nitric Oxide 25, 81–88 (2011).

99. Nzila, A., Ward, S. A., Marsh, K., Sims, P. F. G. & Hyde, J. E. Comparative folate metabolism in humans and malaria parasites (part I): pointers for malaria treatment from cancer chemotherapy. Trends Parasitol. 21, 292–298 (2005).

100. Nzila, A., Ward, S. A., Marsh, K., Sims, P. F. G. & Hyde, J. E. Comparative folate metabolism in humans and malaria parasites (part II): activities as yet untargeted or specific to Plasmodium. Trends Parasitol. 21, 334–339 (2005).

101. Haldar, K., Bhattacharjee, S. & Safeukui, I. Drug resistance in Plasmodium. Nat. Rev. Microbiol. 16, 156–170 (2018).

102. Hyde, J. E. Exploring the folate pathway in Plasmodium falciparum. Acta Trop. 94, 191– 206 (2005).

103. Cosentino, F. & Katusić, Z. S. Tetrahydrobiopterin and dysfunction of endothelial nitric oxide synthase in coronary arteries. Circulation 91, 139–144 (1995).

104. Mitchell, B. M., Dorrance, A. M. & Webb, R. C. GTP cyclohydrolase 1 inhibition attenuates vasodilation and increases blood pressure in rats. Am. J. Physiol. Heart Circ. Physiol. 285, H2165–70 (2003).

105. Pickert, G. et al. Inhibition of GTP cyclohydrolase reduces cancer pain in mice and enhances analgesic effects of morphine. J. Mol. Med. 90, 1473–1486 (2012).

106. Tegeder, I. et al. GTP cyclohydrolase and tetrahydrobiopterin regulate pain sensitivity and persistence. Nat. Med. 12, 1269–1277 (2006).

107. Dai, Y., Cui, J., Gan, P. & Li, W. Downregulation of tetrahydrobiopterin inhibits tumor angiogenesis in BALB/c-nu mice with hepatocellular carcinoma. Oncol. Rep. 36, 669–675 (2016).

108. Chen, L. et al. Paracrine effect of GTP cyclohydrolase and angiopoietin-1 interaction in stromal fibroblasts on tumor Tie2 activation and breast cancer growth. Oncotarget 7, 9353–9367 (2016).

109. Rembold, H. Hemmung des Crithidia-Wachstums durch 4-Amino-pyrimidine. 339, 258– 259 (1964).

110. Xie, L., Smith, J. A. & Gross, S. S. GTP cyclohydrolase I inhibition by the prototypic inhibitor 2, 4-diamino-6-hydroxypyrimidine. Mechanisms and unanticipated role of GTP cyclohydrolase I feedback regulatory protein. J. Biol. Chem. 273, 21091–21098 (1998).

111. Kolinsky, M. A. & Gross, S. S. The mechanism of potent GTP cyclohydrolase I inhibition by 2,4-diamino-6-hydroxypyrimidine: requirement of the GTP cyclohydrolase I feedback regulatory protein. J. Biol. Chem. 279, 40677–40682 (2004).

112. Osei, M. et al. Amplification of GTP-cyclohydrolase 1 gene in Plasmodium falciparum isolates with the quadruple mutant of dihydrofolate reductase and dihydropteroate synthase genes in Ghana. PLoS One 13, e0204871 (2018).

113. Fry, M. & Pudney, M. Site of action of the antimalarial hydroxynaphthoquinone, 2-[trans-4-(4’-chlorophenyl) cyclohexyl]-3-hydroxy-1,4-naphthoquinone (566C80). Biochem. Pharmacol. 43, 1545–1553 (1992).

114. Khanal, S. et al. In Vivo Validation of the Viral Barcoding of Simian Immunodeficiency Virus SIVmac239 and the Development of New Barcoded SIV and Subtype B and C Simian-Human Immunodeficiency Viruses. J. Virol. 94, (2019).

115. Del Prete, G. Q. et al. Molecularly tagged simian immunodeficiency virus SIVmac239 synthetic swarm for tracking independent infection events. J. Virol. 88, 8077–8090 (2014).

116. Weger-Lucarelli, J. et al. Using barcoded Zika virus to assess virus population structure in vitro and in Aedes aegypti mosquitoes. Virology 521, 138–148 (2018).

117. Varble, A. et al. Influenza A virus transmission bottlenecks are defined by infection route and recipient host. Cell Host Microbe 16, 691–700 (2014).

118. Abel, S. et al. Sequence tag-based analysis of microbial population dynamics. Nat. Methods 12, 223–6, 3 p following 226 (2015).

119. Hullahalli, K. & Waldor, M. K. Pathogen clonal expansion underlies multiorgan dissemination and organ-specific outcomes during murine systemic infection. Elife 10, (2021).

120. Fiebig, A. et al. Quantification of Brucella abortus population structure in a natural host. Proc. Natl. Acad. Sci. U. S. A. 118, (2021).

121. Grant, A. J. et al. Modelling within-host spatiotemporal dynamics of invasive bacterial disease. PLoS Biol. 6, e74 (2008).

122. Bachta, K. E. R., Allen, J. P., Cheung, B. H., Chiu, C.-H. & Hauser, A. R. Systemic infection facilitates transmission of Pseudomonas aeruginosa in mice. Nat. Commun. 11, 543 (2020).

123. Pollitt, E. J. G., Szkuta, P. T., Burns, N. & Foster, S. J. Staphylococcus aureus infection dynamics. PLoS Pathog. 14, e1007112 (2018).

124. Zhang, T. et al. Deciphering the landscape of host barriers to Listeria monocytogenes infection. Proc. Natl. Acad. Sci. U. S. A. 114, 6334–6339 (2017).

125. Liu, X. et al. Exploration of Bacterial Bottlenecks and Streptococcus pneumoniae Pathogenesis by CRISPRi-Seq. Cell Host Microbe 29, 107–120.e6 (2021).

126. Wincott, C. J. et al. Cellular barcoding of protozoan pathogens reveals the within-host population dynamics of Toxoplasma gondii host colonization. Cell Rep Methods 2, 100274 (2022).

127. Carrasquilla, M. et al. Barcoding Genetically Distinct Plasmodium falciparum Strains for Comparative Assessment of Fitness and Antimalarial Drug Resistance. MBio 13, e0093722 (2022).

128. Aquilini, E. et al. An Alveolata secretory machinery adapted to parasite host cell invasion. Nat Microbiol 6, 425–434 (2021).

129. Coppens, I. & Joiner, K. A. Host but not parasite cholesterol controls Toxoplasma cell entry by modulating organelle discharge. Mol. Biol. Cell 14, 3804–3820 (2003).

130. Jensen, K. D. C. et al. Toxoplasma polymorphic effectors determine macrophage polarization and intestinal inflammation. Cell Host Microbe 9, 472–483 (2011).

131. Rivera-Cuevas, Y. et al. Toxoplasma gondii exploits the host ESCRT machinery for parasite uptake of host cytosolic proteins. PLoS Pathog. 17, e1010138 (2021).

132. Mayoral, J. et al. Dense Granule Protein GRA64 Interacts with Host Cell ESCRT Proteins during Toxoplasma gondii Infection. MBio 13, e0144222 (2022).

133. Augusto, L. et al. Toxoplasma gondii Co-opts the Unfolded Protein Response To Enhance Migration and Dissemination of Infected Host Cells. MBio 11, (2020).

134. Wang, T. et al. Toxoplasma gondii induce apoptosis of neural stem cells via endoplasmic reticulum stress pathway. Parasitology 141, 988–995 (2014).

135. Zhou, J. et al. Toxoplasma gondii prevalent in China induce weaker apoptosis of neural stem cells C17.2 via endoplasmic reticulum stress (ERS) signaling pathways. Parasit. Vectors 8, 73 (2015).

136. Sugimoto, M. et al. MMMDB: Mouse Multiple Tissue Metabolome Database. Nucleic Acids Res. 40, D809–14 (2012).

137. Jang, C. et al. Metabolite Exchange between Mammalian Organs Quantified in Pigs. Cell Metab. 30, 594–606.e3 (2019).

138. Ma, J. et al. Metabolomic signature of mouse cerebral cortex following Toxoplasma gondii infection. Parasit. Vectors 12, 373 (2019).

139. Zhou, C.-X. et al. Investigation of urine metabolome of BALB/c mouse infected with an avirulent strain of Toxoplasma gondii. Parasit. Vectors 15, 271 (2022).

140. Zhou, C.-X. et al. Metabolomic Profiling of Mice Serum during Toxoplasmosis Progression Using Liquid Chromatography-Mass Spectrometry. Sci. Rep. 6, 19557 (2016).

141. Zhou, C.-X. et al. Serum Metabolic Profiling of Oocyst-Induced Toxoplasma gondii Acute and Chronic Infections in Mice Using Mass-Spectrometry. Front. Microbiol. 8, 2612 (2017).

142. Alday, P. H. & Doggett, J. S. Drugs in development for toxoplasmosis: advances, challenges, and current status. Drug Des. Devel. Ther. 11, 273–293 (2017).

143. Baker, N. et al. Systematic functional analysis of Leishmania protein kinases identifies regulators of differentiation or survival. Nat. Commun. 12, 1244 (2021).

144. Shames, S. R. et al. Multiple *Legionella pneumophila* effector virulence phenotypes revealed through high-throughput analysis of targeted mutant libraries. Proc. Natl. Acad. Sci. U. S. A. 114, E10446–E10454 (2017).

145. Mukhopadhyay, D., Sangaré, L. O., Braun, L., Hakimi, M.-A. & Saeij, J. P. Toxoplasma GRA15 limits parasite growth in IFNγ-activated fibroblasts through TRAF ubiquitin ligases. EMBO J. 39, e103758 (2020).

146. Zamboni, D. S. et al. The Birc1e cytosolic pattern-recognition receptor contributes to the detection and control of Legionella pneumophila infection. Nat. Immunol. 7, 318–325 (2006).

147. Molofsky, A. B. et al. Cytosolic recognition of flagellin by mouse macrophages restricts Legionella pneumophila infection. J. Exp. Med. 203, 1093–1104 (2006).

148. Ren, T., Zamboni, D. S., Roy, C. R., Dietrich, W. F. & Vance, R. E. Flagellin-deficient Legionella mutants evade caspase-1- and Naip5-mediated macrophage immunity. PLoS Pathog. 2, e18 (2006).

149. Diard, M. et al. Stabilization of cooperative virulence by the expression of an avirulent phenotype. Nature 494, 353–356 (2013).

150. Modrzynska, K. et al. A Knockout Screen of ApiAP2 Genes Reveals Networks of Interacting Transcriptional Regulators Controlling the Plasmodium Life Cycle. Cell Host Microbe 21, 11–22 (2017).

151. Lilue, J., Müller, U. B., Steinfeldt, T. & Howard, J. C. Reciprocal virulence and resistance polymorphism in the relationship between Toxoplasma gondii and the house mouse. Elife 2, e01298 (2013).

152. Murillo-León, M. et al. Molecular mechanism for the control of virulent Toxoplasma gondii infections in wild-derived mice. Nat. Commun. 10, 1233 (2019).

153. Dupont, C. D., Christian, D. A. & Hunter, C. A. Immune response and immunopathology during toxoplasmosis. Semin. Immunopathol. 34, 793–813 (2012).

154. Waldman, B. S. et al. Identification of a Master Regulator of Differentiation in Toxoplasma. Cell 180, 359–372.e16 (2020).

155. Sidik, S. M., Huet, D. & Lourido, S. CRISPR-Cas9-based genome-wide screening of Toxoplasma gondii. Nat. Protoc. 13, 307–323 (2018).

156. Markus, B. M., Bell, G. W., Lorenzi, H. A. & Lourido, S. Optimizing Systems for Cas9 Expression in Toxoplasma gondii. mSphere 4, (2019).

157. Hegde, M., Strand, C., Hanna, R. E. & Doench, J. G. Uncoupling of sgRNAs from their associated barcodes during PCR amplification of combinatorial CRISPR screens. PLoS One 13, e0197547 (2018).

158. Prokopenko, D. et al. Utilizing the Jaccard index to reveal population stratification in sequencing data: a simulation study and an application to the 1000 Genomes Project. Bioinformatics 32, 1366–1372 (2016).

159. Cavalli-Sforza, L. L. & Edwards, A. W. Phylogenetic analysis. Models and estimation procedures. Am. J. Hum. Genet. 19, 233–257 (1967).

160. Paradis, E., Claude, J. & Strimmer, K. APE: Analyses of Phylogenetics and Evolution in R language. Bioinformatics 20, 289–290 (2004).

161. Schliep, K. P. phangorn: phylogenetic analysis in R. Bioinformatics 27, 592–593 (2011).

162. Federico, A. & Monti, S. hypeR: an R package for geneset enrichment workflows. Bioinformatics 36, 1307–1308 (2020).

163. Trager, W. & Jensen, J. B. Human malaria parasites in continuous culture. Science 193, 673–675 (1976).

164. Lambros, C. & Vanderberg, J. P. Synchronization of Plasmodium falciparum erythrocytic stages in culture. J. Parasitol. 65, 418–420 (1979).

165. Smith, T. A., Lopez-Perez, G. S., Herneisen, A. L., Shortt, E. & Lourido, S. Screening the Toxoplasma kinome with high-throughput tagging identifies a regulator of invasion and egress. Nat Microbiol 7, 868–881 (2022).

166. Benevides, L. et al. Toxoplasma gondii soluble tachyzoite antigen triggers protective mechanisms against fatal intestinal pathology in oral infection of C57BL/6 mice. PLoS One 8, e75138 (2013).

167. Gazzinelli, R. T., Hakim, F. T., Hieny, S., Shearer, G. M. & Sher, A. Synergistic role of CD4+ and CD8+ T lymphocytes in IFN-gamma production and protective immunity induced by an attenuated Toxoplasma gondii vaccine. J. Immunol. 146, 286–292 (1991).

168. Johnson, J. D. et al. Assessment and continued validation of the malaria SYBR green I-based fluorescence assay for use in malaria drug screening. Antimicrob. Agents Chemother. 51, 1926–1933 (2007).

169. Smilkstein, M., Sriwilaijaroen, N., Kelly, J. X., Wilairat, P. & Riscoe, M. Simple and inexpensive fluorescence-based technique for high-throughput antimalarial drug screening. Antimicrob. Agents Chemother. 48, 1803–1806 (2004).

